# A CRISPR interference system for the nitrogen-fixing bacterium *Azotobacter vinelandii*

**DOI:** 10.1101/2023.11.07.565814

**Authors:** Steven J. Russell, Amanda K. Garcia, Betül Kaçar

## Abstract

A grand challenge for the next century can be found in mitigating the effects of changing climate through bioengineering solutions. Biological nitrogen fixation, the globally consequential, nitrogenase-catalyzed reduction of atmospheric nitrogen to bioavailable ammonia, is a particularly vital area of focus. Nitrogen fixation engineering relies upon extensive understanding of underlying genetics in microbial models, including the broadly utilized gammaproteobacterium, *Azotobacter vinelandii* (*A. vinelandii*). Here we report the first CRISPR interference (CRISPRi) system for targeted gene silencing in *A. vinelandii* that integrates genomically via site-specific transposon insertion. We demonstrate that CRISPRi can repress transcription of an essential nitrogen fixation gene by ∼60%. Further, we show that nitrogenase genes are suitably expressed from the transposon insertion site, indicating that CRISPRi and engineered nitrogen fixation genes can be co-integrated for combinatorial studies of gene expression and engineering. Our established CRISPRi system extends the utility of *A. vinelandii* and will aid efforts to engineer microbial nitrogen fixation for desired purposes.

**For Table of Contents Only:** 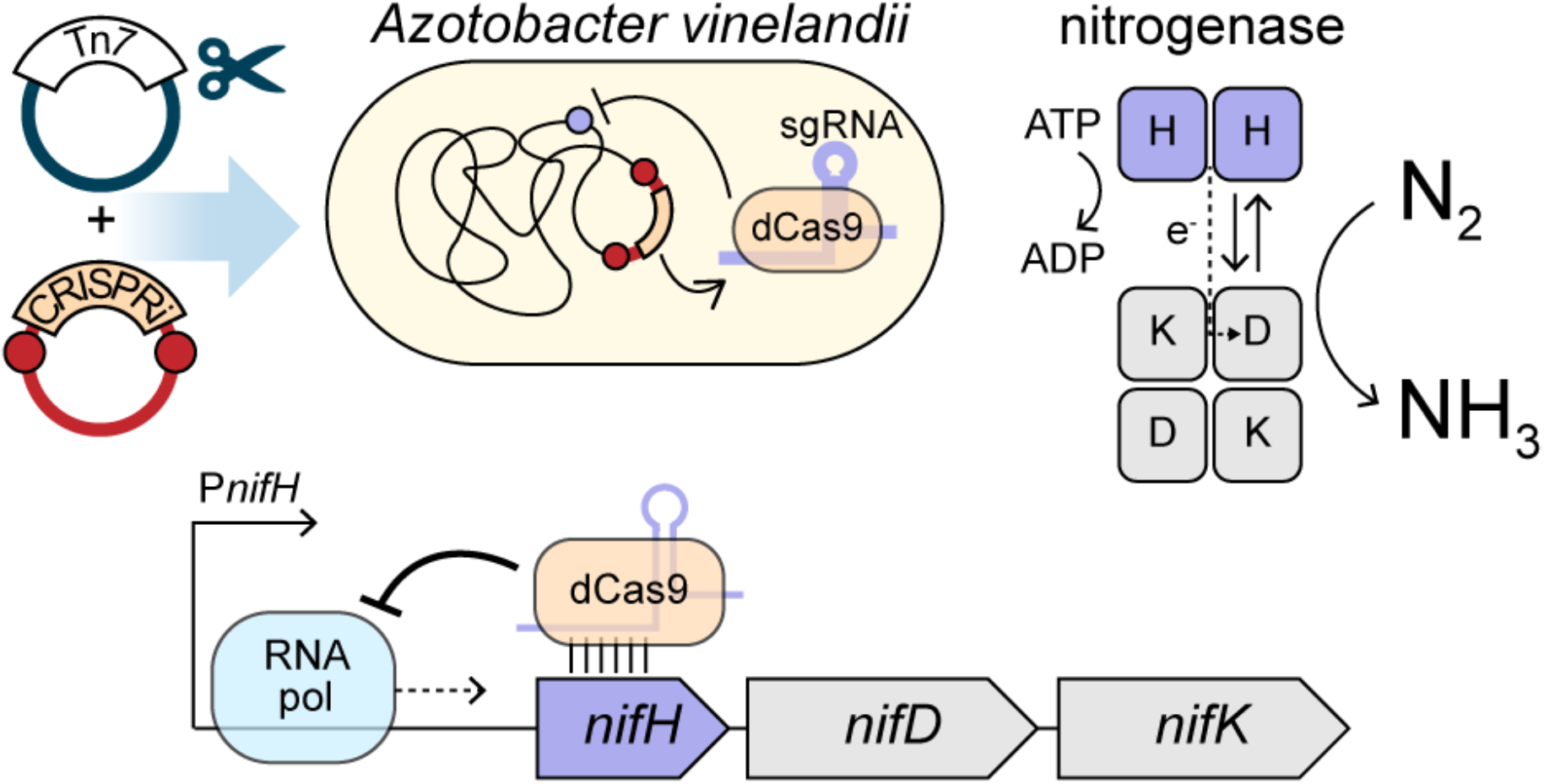

## INTRODUCTION

Biological nitrogen fixation (“N-fixation”, or the enzymatic reduction of atmospheric nitrogen (N_2_) to ammonia (NH_3_), has for >3 billion years been a crucial component of Earth biogeochemical cycling^1–3^. *Azotobacter vinelandii* (*A. vinelandii*), a free-living gammaproteobacterium, is a key model organism for the study and bioengineering of biological N-fixation^4, 5^ (**Figure 1A**). Over the last several decades, *A. vinelandii* has been leveraged to investigate the biochemistry, structure, and evolution of the ancient, core N-fixing enzyme, nitrogenase^6–8^, as well as its supporting biosynthetic and regulatory gene network^9–11^. These principal insights are a prerequisite for N-fixation bioengineering goals, which aim to lessen reliance on the Haber-Bosch process, an industrial method for synthetic N-fixation that today sustains approximately half of the world’s population through crop fertilizer production^12–14^. Despite its role in the agricultural and population boom of the 20^th^ century, the Haber-Bosch process is a leader among chemical manufacturing in nonrenewable energy consumption and greenhouse gas emissions^12,15, 16^. A detailed genetic understanding of N-fixing microbial models like *A. vinelandii* can speed the establishment of alternatives to industrial N-fixation, for example, through the optimization of enzymatic N-fixation regulation and catalysis, improvements to associations between microbial N-fixers and plants, and the transfer of enzymatic N-fixation capability to cereal crops^13, 14, 17, 18^.

**Figure 1.**
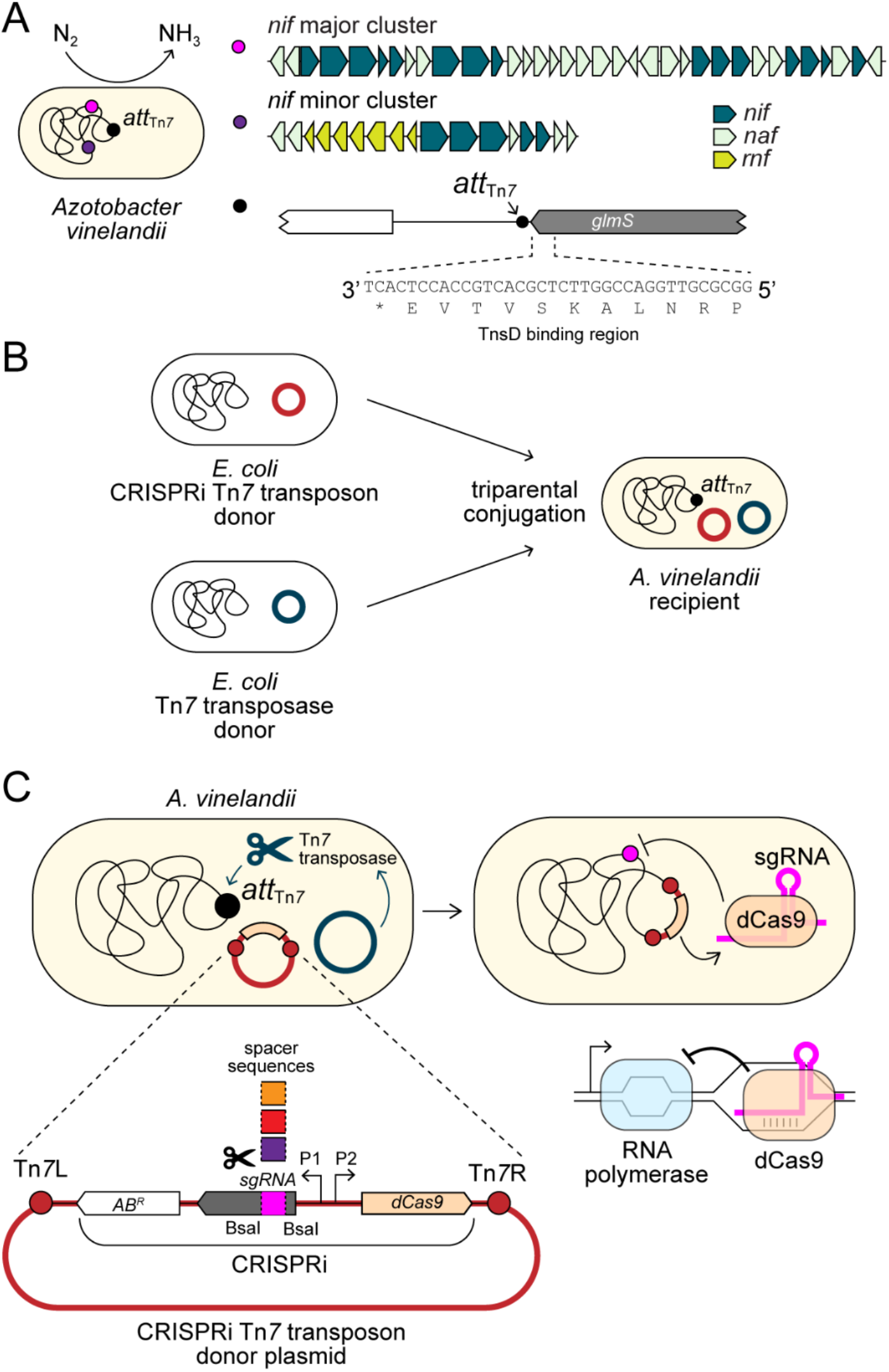
Integration and implementation of a CRISPRi system in *A. vinelandii*. (A) The diazotroph *A. vinelandii* possesses two clusters containing 52 genes (*nif, naf, rnf*) that support assembly and regulation of the molybdenum-dependent nitrogenase enzyme.

A promising tool for the investigation and manipulation of N-fixation genetics in *A. vinelandii* is <u>c</u>lustered <u>r</u>egularly interspaced <u>s</u>hort <u>p</u>alindromic <u>r</u>epeats interference (CRISPRi)^19^. This approach uses an RNA-guided, catalytically deactivated Cas9 nuclease (dCas9) to generate gene knockdowns by sterically blocking transcription of target genes. dCas9 forms a complex with a single guide RNA (sgRNA) and binds to the target gene via base pairing between the sgRNA and DNA. However, the mutant dCas9-sgRNA complex is incapable of cutting the bound DNA due to the loss of nuclease activity, and instead reduces total expression by preventing transcript elongation by RNA polymerase (**Figure 1C**). The phenotypic consequences of gene knockdown can then be systematically assessed and/or leveraged for desired synthetic metabolic systems^20–24^.

Current CRISPRi systems for nitrogen-fixing laboratory models are severely limited^20, 25-^_28_ and, to date, no CRISPRi (nor CRISPR) system has been reported for *A. vinelandii*, despite the organism’s wide applicability to agriculture and biotechnology^5, 14^. Despite important advances generated by gene mutants and knockouts^9, 29–32^, as well as more recently developed tools like transposon sequencing^33^, the functional roles and essentiality of many N-fixation-related genes under different physiological conditions have yet to be fully characterized^4^. Further, the *A. vinelandii* model lacks the associated genetic tools for the modular manipulation of N-fixation and N-fixation-related gene expression, which would otherwise make them amenable to optimization under desired conditions.

Here, we report a CRISPRi system for *A. vinelandii*, the first CRISPR-based genetic tool for this widely used bacterial N-fixation model. We demonstrate the successful transfer of CRISPRi machinery into *A. vinelandii* using Mobile-CRISPRi^34^ vectors and Tn*7*-transposon-based genomic integration. CRISPRi components are stably integrated into the *A. vinelandii* genome and result in ∼50% knockdown of fluorescence reporters co-integrated at the Tn*7* insertion site. Further, we report constructs to perform knockdown of native nitrogenase structural genes, achieving ∼60% transcription suppression of the essential N-fixation electron transfer protein, NifH, under diazotrophic growth conditions. Finally, we confirm that the Tn*7* insertion site is a suitable locus for expression of structural nitrogenase genes, permitting simultaneous genomic integration and investigation of CRISPRi and engineered N-fixation genetic components in *A. vinelandii*.

The *A. vinelandii* genome also contains a 5’ *glmS* sequence motif that is highly conserved among gammaproteobacteria and serves as the TnsD binding region for downstream Tn*7* transposon insertion at *att*_Tn*7*_. (B) Mobile-CRISPRi plasmids are transferred to *A. vinelandii* via triparental conjugation. Two *E. coli* donor strains harbor plasmids with either the Tn*7* transposon containing the CRISPRi machinery (red) or the Tn*7* transposase machinery (dark teal). (C) The Tn*7* transposase enables genomic insertion of the CRISPRi transposon at the *A. vinelandii att*_Tn*7*_ site. Generalized schematic of the CRISPRi Tn*7* transposon is shown (lower left). Expression of dCas9 and sgRNA is controlled by separate inducible promoters of choice, P1 and P2 (P_L*lacO1*_ used in the present study; see Figure 2). Unique spacer sequences are cloned into the Tn*7* transposon to program knockdown of target genes within the *A. vinelandii* genome. The expressed dCas9 and sgRNA complex binds to the target region via base pairing between genomic DNA and the unique spacer sequence, interfering with transcript elongation by RNA polymerase (lower right).

**Figure 2.**
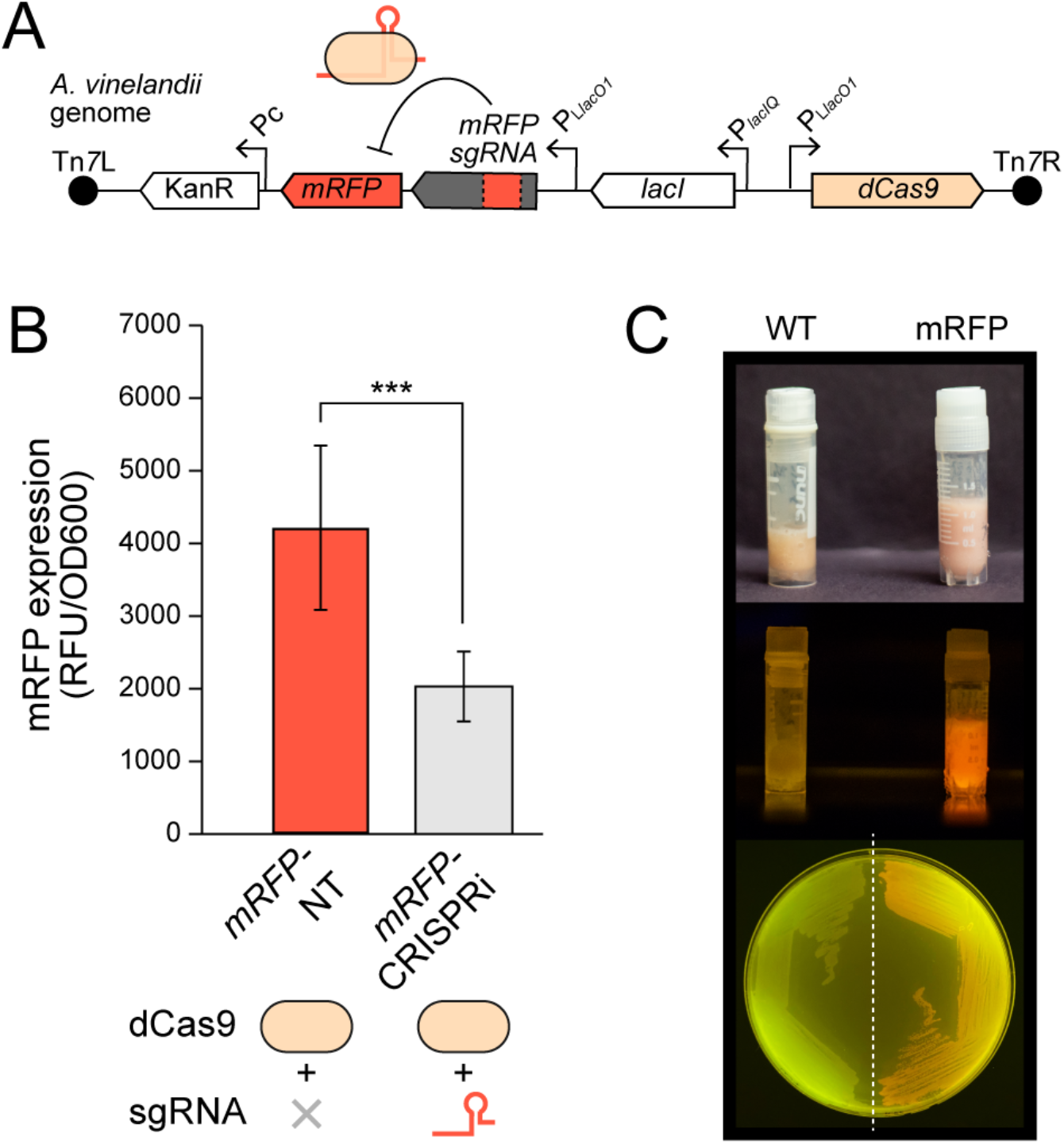
CRISPRi knockdown of a genomically integrated fluorescence reporter gene in *A. vinelandii*. (A) Schematic of the Tn*7* transposon inserted into the *A. vinelandii* genome, which contains *dCas9*, *mRFP*, and a sgRNA cassette with spacer sequence complementary to *mRFP*. (B) Knockdown of mRFP expression in *A. vinelandii* cells assessed by fluorimetry. Fluorescence values are normalized to cell density (OD600) and corrected against baseline WT fluorescence (lacking *mRFP*). Each bar in the plot represents the mean of nine biological replicates with error bars indicating ± 1 SD. ***: *p* < 0.001. (C) Images of *A. vinelandii* mRFP fluorescent phenotype from both strain stocks (top under natural light, middle under 465-nm blue light) and plated cells (bottom, under 465-nm blue light). Nonfluorescent WT strain shown for comparison.

## RESULTS AND DISCUSSION

### CRISPRi machinery integrates into the *A. vinelandii* genome and achieves knockdown of target genes

To develop a CRISPRi system for *A. vinelandii*, we leveraged the Mobile-CRISPRi system, a suite of modular vectors for the conjugative transfer and genomic integration of dCas9 machinery into diverse bacteria^34^ (Figure 1). Development of CRISPRi for new hosts can be challenged by limitations to existing genetic tools for its transfer and incorporation, potential toxicity of expressed dCas9 or sgRNA (the latter resulting from off-target effects ^35^), and poor knockdown efficiencies, all requiring testing and, if needed, additional optimization^21, 36^. Mobile-CRISPRi, which includes components based on the *Streptococcus pyogenes* (*Spy*) dCas9 enzyme, broadens the transferability of CRISPRi to proteobacteria by leveraging the Tn*7* transposase system for genomic integration. Further, the modularity of its associated vectors enables improved specificity and optimization of CRISPRi parts. Mobile-CRISPRi was previously used to transfer CRISPRi components into several model (e.g., *Staphylococcus aureus*, *Listeria monocytogenes*, *Pseudomonas aeruginosa*, *Zymomonas mobilis*) and non-model (e.g., *Vibrio casei*) proteobacteria and Firmicutes^21, 34^. Thus, this approach is promising for the development of a CRISPRi system in the gammaproteobacterium *A. vinelandii*.

We first assessed the feasibility of Tn*7* integration of CRISPRi machinery by identifying the Tn*7* transposon attachment site, *att*_Tn*7*_, in *A. vinelandii*, downstream of the *glmS* gene (Figure 1A)^37^. We confirmed the presence of a single *att*_Tn*7*_ site on the *A. vinelandii* chromosome. The 3’ end of *glmS* contains a sequence motif highly conserved across gammaproteobacteria that serves as a binding site for the transposase target selector protein TnsD^38^. Further, the 3’ end of *glmS* in *A. vinelandii* is >500 bp from the nearest open reading frame, indicating that Tn*7* insertion would not likely disrupt downstream gene function and that *att*_Tn*7*_ would thus act as a neutral insertion site for CRISPRi components^39, 40^.

We attempted CRISPRi transfer to *A. vinelandii* using Mobile-CRISPRi test constructs that contain both CRISPRi as well as a fluorescence reporter cassette for initial determination of gene knockdown efficiency. These components include genes encoding *Spy* dCas9, monomeric red fluorescent protein (mRFP), sgRNA targeting *mRFP*, and a kanamycin antibiotic resistance marker (KanR) for Tn*7* insertion selection^36^ (Figure 2A). Both sgRNA and dCas9 expression are controlled by IPTG-inducible P_L*lacO1*_ promoters. Wild-type (WT) recipient *A. vinelandii* cells were mated with two *Escherichia coli* (*E. coli*) donor strains, one carrying a CRISPRi Tn*7* transposon vector and the other carrying a Tn*7* transposase vector (Figure 1B). We isolated two *A. vinelandii* transconjugants, a “targeting” strain that possesses both *dCas9* and *mRFP*-targeting sgRNA (strain “*mRFP*-CRISPRi”) and a “non-targeting” strain with *dCas9* but lacking a sgRNA cassette (strain “*mRFP*-NT”) (**Table 1**).

**Table 1.**
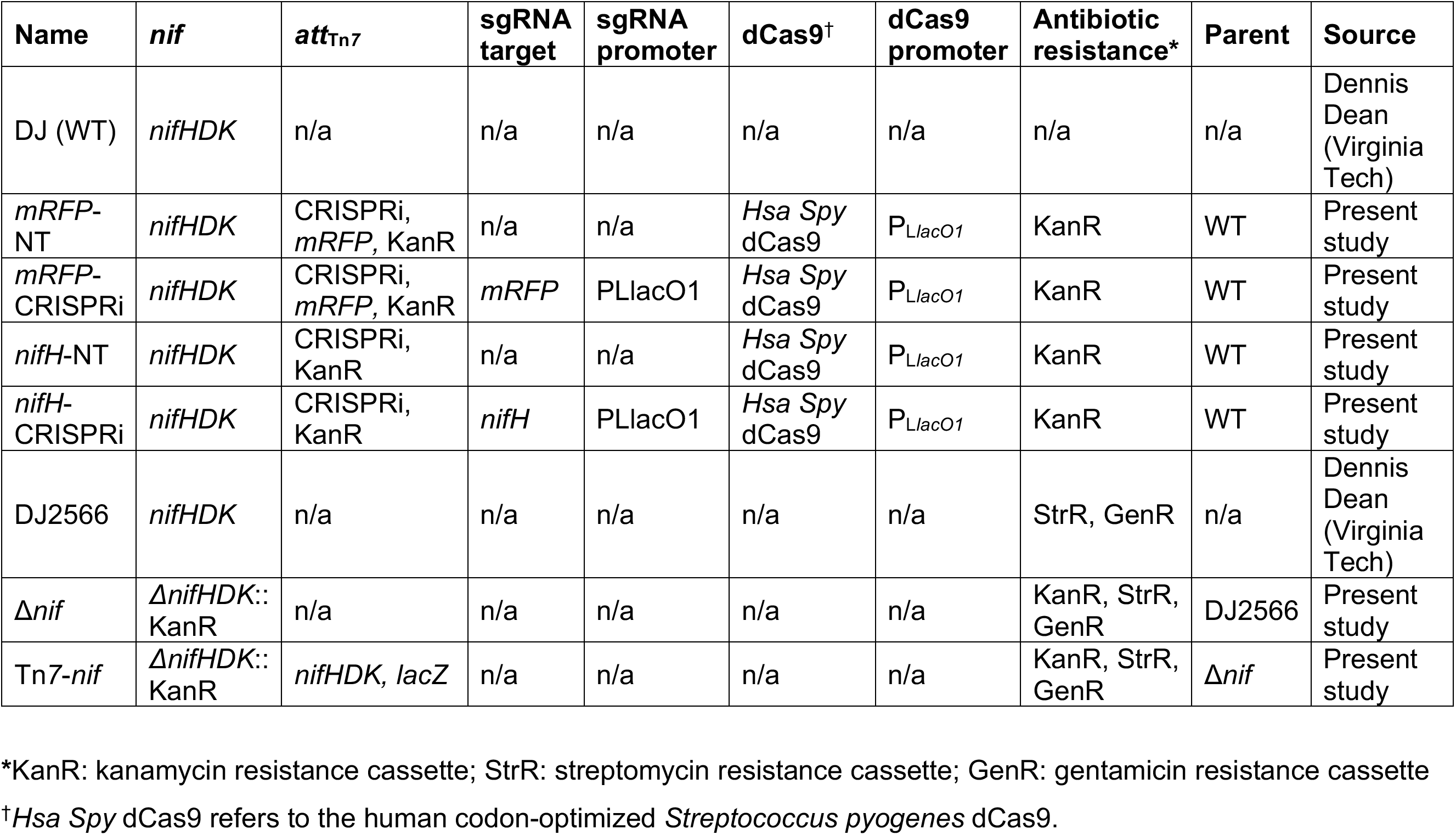
Summary of *A. vinelandii* strains used and constructed in the present study (see **Table S1** for additional details).

Gene knockdown efficiency was measured by fluorimetry of *A. vinelandii* cells following induction of the CRISPRi system targeting the co-integrated *mRFP* fluorescence reporter. Strain *mRFP*-CRISPRi cultured with IPTG (carrying both dCas9 and the *mRFP*-targeting sgRNA) showed an average 52% reduction relative to the non-targeting *mRFP*-NT strain (*p* = 0.0001) (Figure 2B). Our results thus demonstrate successful induction of both dCas9 and sgRNA as well as repression of the *mRFP* target. Further, growth of both targeting and non-targeting strains establish that neither dCas9 nor sgRNA expressed at the tested levels is toxic to *A. vinelandii*.

### CRISPRi knockdown reduces expression of proteins essential for N-fixation

We next tested CRISPRi knockdown efficiency of native nitrogen fixation genes encoding the nitrogenase enzyme in *A. vinelandii*. We designed sgRNA cassettes targeting the *nifHDKTY* operon, the latter which is under control of the *nifH* promoter (P*_nifH_*) and is expressed in the absence of a fixed nitrogen source in the culture medium^10, 11, 41^ (Figure 3A). This operon includes genes that code for the molybdenum-dependent nitrogenase enzyme complex (NifHDK), as well as the accessory nitrogen fixation proteins NifT and NifY, whose disruption was previously determined to have moderate or weak negative fitness effects *A. vinelandii*^33^. Under diazotrophic conditions, nitrogenase proteins are among the most highly expressed in *A. vinelandii*^10^ – comprising ∼10% of total protein^42^ – creating a broad dynamic expression range for CRISPRi engineering.

**Figure 3.**
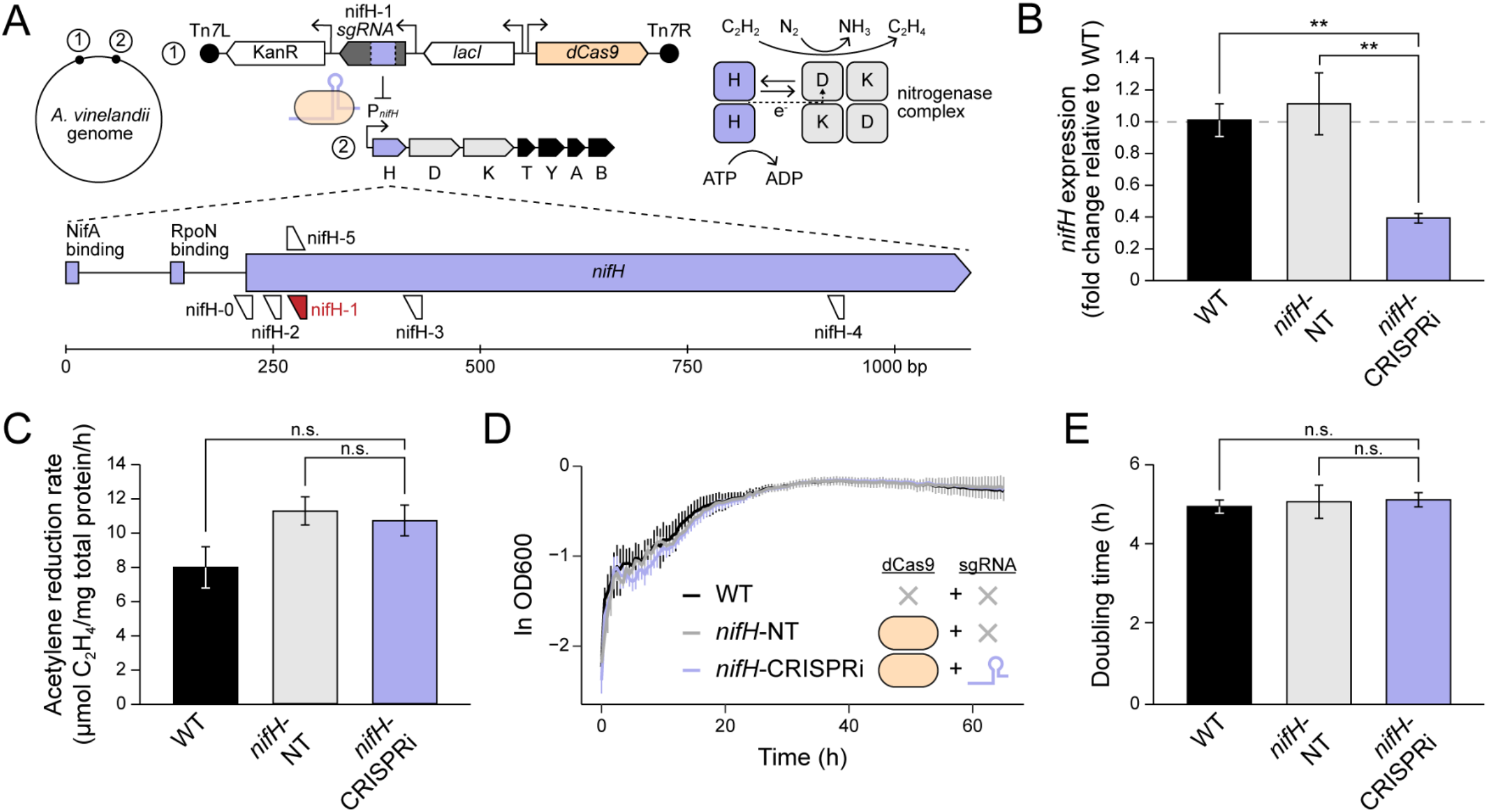
CRISPRi knockdown of native nitrogenase in *A. vinelandii*. (A) Schematic of the Tn*7* transposon inserted into the *A. vinelandii* genome containing *nifH*-targeting CRISPRi machinery. The NifH protein, an essential component of nitrogenase, transfers electrons to the rest of the enzyme complex, ultimately resulting in the reduction of N_2_ to NH_3_ (as well as the alternate substrate acetylene (C_2_H_2_) to ethylene (C_2_H_4_)). Five sgRNA spacer sequences (nifH-0 to nifH-5) complementary to genomic DNA between the P*_nifH_* promoter and the 3’ end of *nifH* were cloned into Tn*7* transposon plasmids. The construct containing sgRNA nifH-1 (red) was successfully transferred to *A. vinelandii*. (B) CRISPRi knockdown of *nifH* expression assessed by RT-qPCR. (C) Acetylene reduction rates of *A. vinelandii strains* following nitrogenase derepression. (D) Diazotrophic growth curve of *A. vinelandii* strains. Curves are averaged from three biological replicates per strain and error bars indicate ± 1 SD. (E) Doubling times of *A. vinelandii* strains during diazotrophic growth. (B, D, E) Each bar represents the mean of 3 biological replicates, with the exception of that for WT in (B) and (C), which represents the mean of two biological replicates. Error bars indicate ± 1 SD. **: *p* < 0.01; n.s.: not significant, *p* > 0.05.

We designed five sgRNA cassettes, each containing spacer sequences complementary to sense and antisense DNA strands located between promoter P*_nifH_* and the *nifH* 3’ end, the first gene within the *nifHDKTY* operon (Figure 3A, **Table 2**). We designed multiple sgRNA sequences due to the previously demonstrated sensitivity of knockdown efficiency to the target binding region. For example, knockdown efficiency can vary depending on whether the spacer sequence is complementary to sense or antisense DNA strands^19, 43^, and can also vary according to the spacer binding position across the length of the promoter and coding region^44, 45^. Tn*7* transposon vectors, each cloned with one sgRNA cassette under the PLlacO1 promoter, were conjugated to WT *A. vinelandii* recipient cells. One construct, containing sgRNA “nifH-1”, targeting the antisense strand with +52-bp offset from the *nifH* start codon, was successfully transferred and inserted into *A. vinelandii*, yielding strain “*nifH*-CRISPRi”. A corresponding non-targeting strain, lacking an sgRNA cassette, was also isolated (strain “*nifH*-NT”).

**Table 2.**
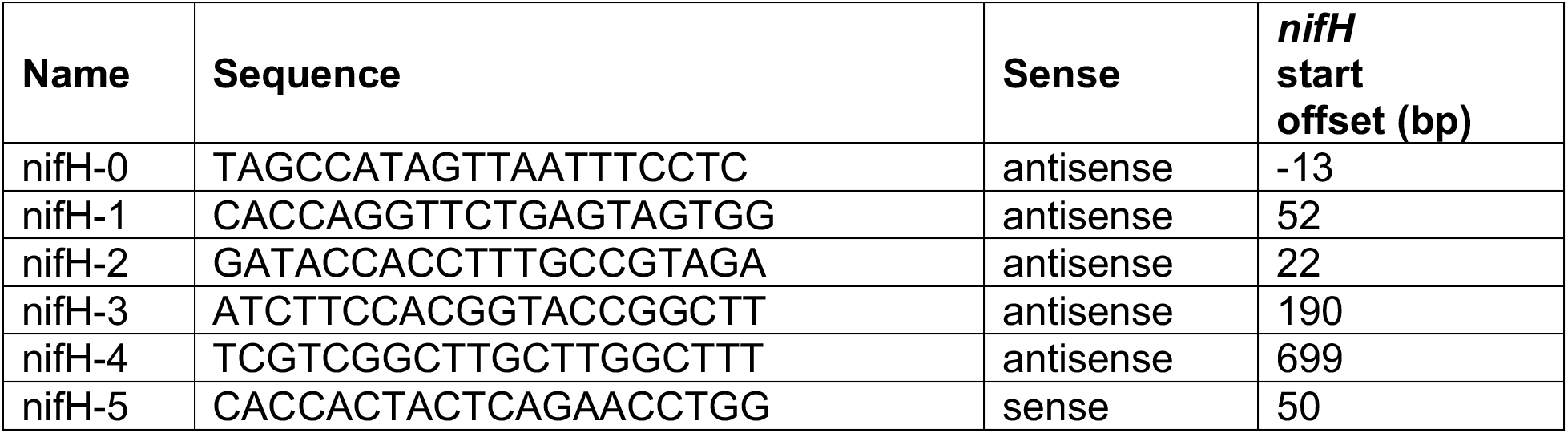
sgRNA sequences targeting knockdown of the *nifHDKTY* operon. Sequences bind sense or anti-sense DNA upstream or within the *nifH* gene.

Expression knockdown of the *nifHDKTY* operon in strain *nifH*-CRISPRi was assessed by measuring *nifH* transcript levels by RT-qPCR, following both induction of the CRISPRi system as well as derepression of nitrogenase (see Methods). The depression approach permitted the culturing of sufficient cell densities with an exogeneous fixed N source prior to their transfer to diazotrophic conditions (i.e., lacking fixed N) and subsequent activation of the *nifHDKTY* operon. We detected a ∼60% decrease in *nifH* transcript in strain *nifH*-CRISPRi relative to both WT *A. vinelandii* (*p* = 0.009) and non-targeting *nifH*-NT strains (*p* = 0.003) (Figure 3B). Our results demonstrate successful repression of *nifH* by CRISPRi.

We further measured the phenotypic impact of *nifH* knockdown by characterizing diazotrophic growth and acetylene reduction activity following CRISPRi induction. Reduction of acetylene, an alternative substrate of the nitrogenase, to ethylene classically serves as a proxy for *in vivo* nitrogenase activity in *A. vinelandii*^46^. Despite the ∼60% decrease in *nifH* transcript following nitrogenase derepression, acetylene reduction rates of the same *nifH*-CRISPRi samples were comparable between WT (*p* = 0.055) and *nifH*-NT strains (*p* = 0.75) (Figure 3C). Doubling times of all *A. vinelandii* strains under standard diazotrophic conditions were similarly comparable (*p* = 0.48; Figure 3D**, 3E**).

### Neutral expression of nitrogenase genes at the Tn*7* insertion site

After confirming CRISPRi knockdown of native nitrogenase genes, we designed a construct to expand on the utility of CRISPRi for investigations of nitrogen fixation genetics by expressing nitrogenase genes from the Tn*7* insertion site. The Tn*7* site is typically neutral for insertion (i.e., does not result in a detectable phenotypic impact)^40^, provided that the *att*_Tn*7*_ site does not disrupt neighboring genes^39^, which we have not found to be the case in *A. vinelandii* (see above; Figure 1A). Confirmation that nitrogenase genes can be successfully expressed from the Tn*7* site would permit the simultaneous incorporation of CRISPRi components targeting N-fixation genes as well as engineered or refactored N-fixation genes via Tn*7* insertion.

To test nitrogenase expression from the Tn*7* insertion site, a nitrogenase deletion strain, “Δ*nif*”, of *A. vinelandii* was first constructed, containing disruptions in all three *nif*, *vnf*, and *anf* gene clusters, the latter two encoding the alternative V-and Fe-only-dependent nitrogenases that is expressed under Mo-starvation^11^ (**Table 1**). We then relocated *nifHDK* by inserting these genes back into the *A. vinelandii* chromosome by Tn*7* transposase at the *att*_Tn*7*_ site, generating strain “Tn*7*-*nif*” (Figure 4A). Tn*7*-*nif* grew comparably to WT *A. vinelandii* under Mo-replete, diazotrophic conditions, with no significant difference in doubling time (*p* = 0.33) albeit a ∼13% increase in growth lag time (*p =* 0.009) (Figure 4B**-4D**). By contrast, the Δ*nif* deletion strain from which Tn*7*-*nif* was constructed exhibited severe defects in diazotrophic growth. These results demonstrate that the reintroduction of *nifHDK* genes at the *att*_Tn*7*_ site of Δ*nif* is sufficient to rescue diazotrophic growth, and that expression of nitrogenase protein from *att*_Tn*7*_ is suitable for downstream N-fixation studies in *A. vinelandii*, including future co-integration of CRISPRi and nitrogenase components.

**Figure 4.**
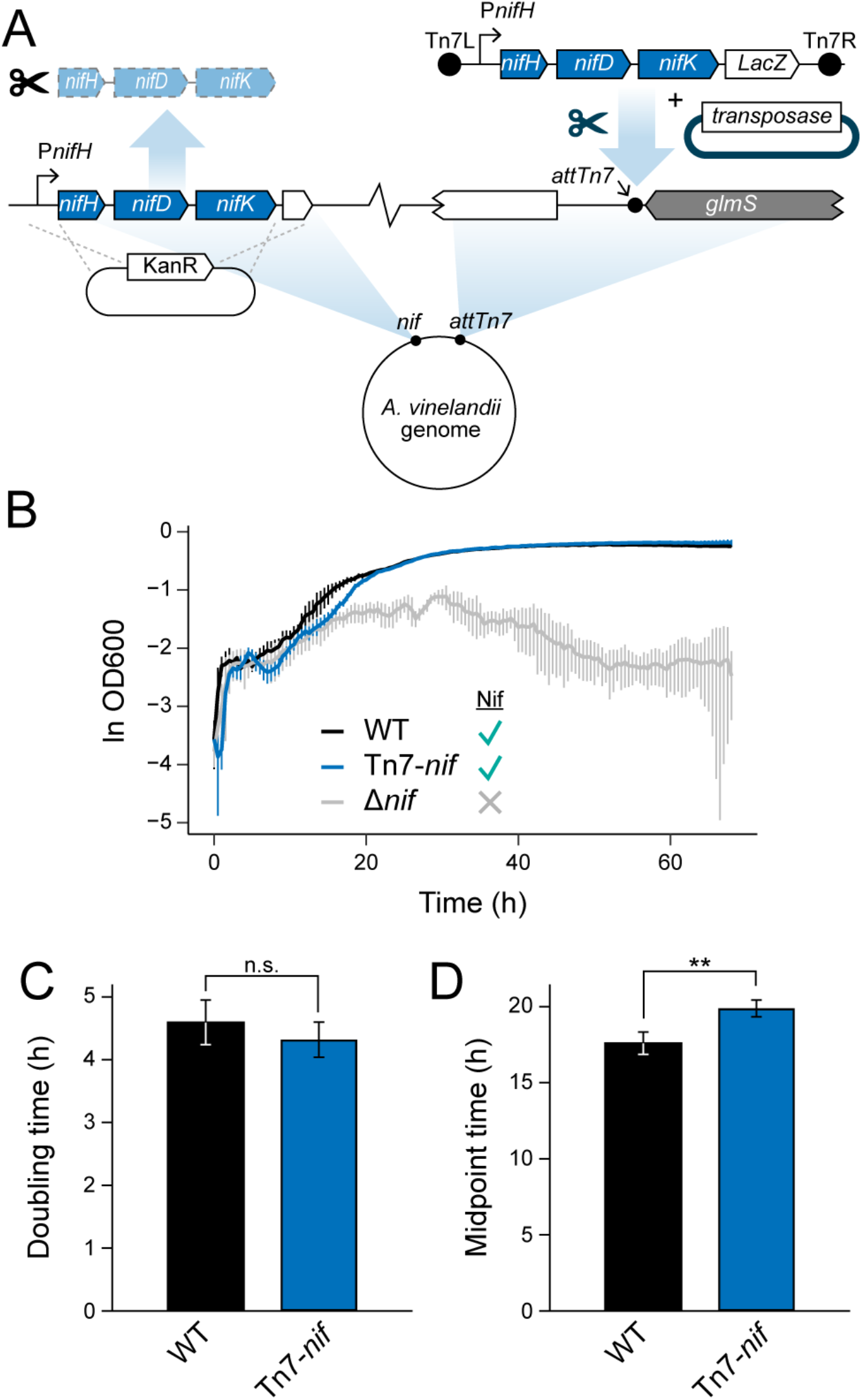
Neutral expression of nitrogenase *nifHDK* genes from the *A. vinelandii att*_Tn*7*_ site. (A) Relocation of native *nifHDK* genes to *att*_Tn*7*_ via Tn*7* transposon insertion. Native *nifHDK* was cleanly excised and replaced with a KanR cassette via homologous recombination. WT *nifHDK* genes under the P*_nifH_* promoter were cloned into a Tn*7* transposon and genomically integrated into the Δnif strain with Tn*7* transposase. (B) Diazotrophic growth curve of *A. vinelandii* strains. Curves are averaged from three biological replicates per strain and error bars indicate ± 1 SD. (C) Doubling times of *A. vinelandii* strains during diazotrophic, exponential growth. (D) Midpoint times of *A. vinelandii* strains, representing the inflection point of a logistic model fit to the growth curve. (C,D) Each bar represents the mean of 3 biological replicates and error bars indicate ± 1 SD. **: *p* < 0.01; n.s.: not significant, *p* > 0.05.

## Discussion

Relative to known diazotrophs, *A. vinelandii* possesses a highly complex genetic system for N-fixation, hosting 52 genes within clusters that support the Mo-dependent nitrogenase system (Nif), as well as additional genes that support the alternative V-(Vnf)and Fe-only (Anf) nitrogenases^4, 47^. By comparison, some diazotrophs, namely certain thermophilic archaea, can perform N-fixation with less than ten genes with known counterparts in *A. vinelandii*^48^. On one hand, this genetic complexity has been advantageous for N-fixation studies, providing source material for a wealth of scientific literature on nitrogenase enzymology^6, 7, 49^, metallocluster assembly^9^, regulation in response to fixed nitrogen source, metal availability, and nitrogenase mutations^10, 11, 41^, electron transport to support the energy intensive N-fixation pathway^50, 51^, and coordination of the three variably metal-dependent nitrogenase systems^52, 53^, including the Mo-dependent Nif nitrogenase studied here. In principle, these insights might be extended to other diazotrophs which contain subsets of N-fixation genes present in *A. vinelandii*, with the assumption that fewer genes represent a simplification of genetic schemes possessed by *A. vinelandii*, rather than altogether different schemes. Additionally, it has been suggested that the genetic complexity of diazotrophs like *A. vinelandii* have resulted from the optimization of N-fixation systems for an obligately aerobic bacterium^47^. Thus, insights from *A. vinelandii* genetics are invaluable for understanding what is required to assemble and maintain the oxygen-sensitive nitrogenase in aerobic organisms, a significant hurdle for the goal of transferring N-fixation capability to cereal crops^13, 14^.

Nevertheless, the numerous N-fixation genes in *A. vinelandii* also pose significant experimental challenges for their functional interrogation and use in bioengineering. With so many targets, traditional genetic approaches for introducing mutations and gene deletions (via homology-directed recombination) are impractical for high-throughput analyses. Further, the complexity of *A. vinelandii* genetics is reflected in oftentimes recalcitrant regulatory schemes for its N-fixation components. For example, study of the alternative V-and Fe-only-dependent nitrogenase systems in *A. vinelandii* can be hindered by their repression given even trace abundances of free Mo^30^. Finally, such complexity challenges facile predictions for how the expansive N-fixation network would respond to sequence-level engineering^8, 10^, metabolic refactoring^54, 55^, and differing environmental conditions of *A. vinelandii* or other diazotrophs with similar N-fixation genetics.

CRISPRi (and, particularly, the Mobile-CRISPRi implementation leveraged in the present study) poses several advantages over other genetic manipulation strategies that have previously been used to probe N-fixation gene function and essentiality in *A. vinelandii*. Both gene knockouts and transposon sequencing, the latter recently leveraged to quantify gene fitness effects in *A. vinelandii*^33^, are effective for identifying genes that contribute significantly to host fitness, but do not permit subsequent analysis of the phenotypic response if gene disruption proves lethal. Transposon sequencing avoids some of the disadvantages of traditional gene disruption methods by its high-throughput nature and is additionally not limited by effective selection markers. However, high sequencing coverage and the creation of large libraries are required^56^. CRISPRi, like transposon sequencing, can permit the high-throughput characterization of N-fixation genes in *A. vinelandii* and other diazotrophs via the creation of sgRNA spacer sequences that enable targeted silencing of genes throughout the genome. Importantly, however, CRISPRi can be used to generate partial knockdowns and can be made to be inducible, titratable, and reversible^19, 34^, allowing further characterization of essential gene response to varying levels of expression, and can be multiplexed to repress several genes simultaneously^57^.

A significant benefit of the Mobile-CRISPRi vector suite used in the present study is the modularity of its components, including the dCas9 or sgRNA sequences as well as each of their promoters, which permits greater optimization potential. Previous work has shown that CRISPRi knockdown efficiency can be controlled through placement of *dCas9* and *sgRNA* genes under inducible promoters, and the extent of gene knockdown can be manipulated by alteration of promoter strength^58^, truncation of spacer sequences^59^, or introduction of base-pair mismatches between the sgRNA and the target gene^60^. The knockdown efficiencies for the *A. vinelandii* CRISPRi system reported here could be optimized by incorporating these strategies, for example by using more effective promoters for *A. vinelandii,* including the strong, native nitrogenase P*_nifH_* promoter itself. Further, larger libraries of spacer sequences could be constructed to identify sgRNA cassettes for optimized N-fixation gene silencing. We expect future work to expand on the utility of the CRISPRi system established here through such strategies and broader application to the *A. vinelandii* N-fixation network.

## CONCLUSIONS

In this study, we developed a CRISPRi system for *A.* vinelandii, the first CRISPR system reported for the widely utilized N-fixing model gammaproteobacterium. We show that the designed CRISPRi components achieve 50-60% targeted knockdown of both native and non-native genes, including those essential for the expression of the core N-fixing enzyme, nitrogenase. The *A. vinelandii* CRISPRi system established here, together with sequence-level manipulations of N-fixation components, provides an ideal groundwork for future, combinatorial investigations of N-fixation genetics, evolution, and bioengineering. In particular, such approaches could be used to tune N-fixation genetic network response to phylogenetically guided, protein-sequence-level engineering of nitrogenase proteins, which would expand understanding and utility of nitrogenase catalytic diversity, both past and present^8, 10^.

## METHODS

### Bacterial strains and culture conditions

All *A. vinelandii* and *E. coli* strains used or generated in the present study are described in **Table 1 and** **Table S1**. *A. vinelandii* strains are derived from the DJ strain (ATCC BAA-1303; referred to here as wild-type, “WT”) generously provided by Dennis Dean (Virginia Tech).

*A. vinelandii* cells were cultured in liquid or solid Burk’s media (“B medium”) containing 1 µM Na_2_MoO_4_, with 10 mM ammonium acetate (yielding “BN medium”), 1 mM IPTG, 0.6 µg/mL kanamycin, 100 µg/mL X-Gal, and/or yeast extract (Fisher Scientific Cat. No. BP9727; 5 g added per L of growth medium) amendments as needed (see below). Cells were grown at 30 °C and liquid cultures were shaken orbitally at 300 rpm.

*E. coli* cells were cultured in liquid or solid Luria-Bertani medium (“LB medium”; Sigma Aldrich Cat. No. L3522), with 100 µg/mL carbenicillin and 300 µM diaminopimelic acid (DAP) amendments as needed (see below). Cells were grown at 37 °C and liquid cultures were shaken orbitally at 250 rpm.

### Plasmid construction

Plasmids in the present study were obtained or modified from the Mobile-CRISPRi suite^34,36^ and are described in **Table S2**. Primers and synthetic oligonucleotides used for plasmid construction are listed in **Table S3.** pJMP1187 and pJMP1189 plasmids for testing Tn*7*-insertion of CRISPRi components and knockdown of the mRFP reporter in *A. vinelandii* are described previously and were generously donated by Jason Peters (University of Wisconsin-Madison)^34, 36^.

Construction of plasmids targeting *nifH* knockdown followed Banta et al.^36^ with modifications as described below. A sgRNA spacer sequence library for the *A. vinelandii* genome was generated via the build_sgrna_library.py code obtained from https://github.com/ryandward/sgrna_design. Five spacer sequences complementary to the genomic region between P*_nifH_* and the 3’ end of *nifH* were selected (nifH-0 to nifH-5). Single-stranded spacer oligonucleotides and their complements were synthesized with 4-bp BsaI overhangs (Integrated DNA Technologies, Coralville, IA, USA) and subsequently annealed. The resulting double-stranded DNA fragments were individually cloned into pJMP1339 following digestion with BsaI (New England Biolabs Cat. No. R3733), purification (Monarch PCR & DNA Cleanup Kit, New England Biolabs Cat. No. T1030), and ligation with T4 DNA ligase (New England Biolabs Cat. No. M0202), generating plasmids pSR38-pSR43 (see **Figure S1** for plasmid map of pSR39, containing sgRNA-nifH-1). A non-targeting plasmid, pSR44, was constructed by digestion of pJMP1339 with EcoRI to excise the sgRNA cassette followed by self-ligation (see **Figure S2** for plasmid map).

Plasmid pSR37 for Tn*7*-based insertion of *nifHDK* genes at the *A. vinelandii att*_Tn*7*_ site was constructed by NEBuilder HiFi assembly (New England Biolabs Cat. No. E2621) from the pJMP6957 Tn*7* vector, lacking both CRISPRi and sgRNA cassettes (see **Figure S3** for plasmid map). A synthetic *lacZ* cassette (Twist Biosciences, South San Francisco, CA, USA) and a 4,653-bp *A. vinelandii* gDNA fragment containing *nifHDK* was PCR-amplified with primers 533/534 and 535/536, respectively, containing appropriate homologous sequences for HiFi assembly. pJMP6957 was digested with XhoI (New England Biolabs Cat. No. R0146) and purified (QIAquick PCR Purification Kit, Qiagen Cat. No. 28104) to excise the spectinomycin resistance cassette. The *lacZ* cassette was cloned into the pJM6957 plasmid via NEBuilder HiFi assembly, yielding plasmid pSR36. pSR36 was then digested with PmeI (New England Biolabs Cat. No. R0560) to excise the *sfGFP* cassette and assembled with the *nifHDK* fragment via NEBuilder HiFi assembly, generating plasmid pSR37.

All plasmid constructs were confirmed by whole-plasmid Oxford Nanopore sequencing (Plasmidsaurus, Eugene, OR, USA). Sequence-confirmed plasmids were transformed into *E. coli* mating strain WM6026 by electroporation for subsequent conjugation of *A. vinelandii* (see below).

### *A. vinelandii* strain construction

*A. vinelandii* mutant strains were constructed via Tn*7*-based genomic insertion of CRISPRi, antibiotic resistance marker, fluorescence reporter, and/or nitrogenase genetic components. Tn*7* transposon or transposase plasmids, hosted by *E. coli* mating strains (derivatives of WM6026), were transferred to *A. vinelandii* via tri-parental conjugation following Banta et al.^36^, with modifications as described below. *A. vinelandii* parent cells were cultured in 50 mL BN medium to saturation (20-24 h). *E. coli* mating cells from two strains, the first harboring the desired Tn*7* transposon plasmid and the second, sJMP2954, containing the Tn*7* transposase plasmid pJMP1039, were cultured in 5 mL LB medium + DAP to saturation (16-18 h). 100 µL of each of the three strains was combined with 700 µL BN medium with yeast extract (“BNY medium”), followed by centrifugation at 7,000 x g for 2 min. The resulting cell pellet was resuspended in 30 µL BNY medium, transferred to a nitrocellulose filter on solid BNY medium plates, and incubated at 37 °C for 24 h. The nitrocellulose filter was then transferred to a 1.5 mL centrifuge tube and suspended in 200 µL 1x PBS, pH 7.4. Suspension aliquots were plated on appropriate selective solid medium: BN medium + X-Gal for strain Tn*7*-*nif* and BN medium + kanamycin for all other *A. vinelandii* transconjugants. Plated cells were incubated at 30 °C for 2-4 days. Transconjugants were screened by repeated scoring on selective solid medium and Tn*7* insertion was confirmed by PCR amplification of the *att*_Tn*7*_ site and Sanger sequencing with appropriate primers (**Table S3**; **Figure S4**). All *A. vinelandii* transconjugants were additionally confirmed by Oxford Nanopore whole-genome sequencing (Plasmidsaurus, Eugene, OR, USA) (NCBI number pending).

*A. vinelandii* strain Δ*nif*, the parent of strain Tn*7*-*nif*, contains disruptions to all three native nitrogenase gene clusters (*nif, vnf, anf*) and was constructed from DJ2566 (inactivated for *vnf* and *anf* only; generously donated by Dennis Dean, Virginia Tech), following Dos Santos et al.^61^. Competent DJ2566 cells were prepared via metal starvation (i.e., growth in BN medium lacking both Mo and Fe salts). Competent cells were transformed with ∼1000 ng of plasmid pAG25 containing a KanR cassette and flanking sequences directing replacement of *nifHDK* genes by homologous recombination. Cells were plated on solid BN medium + kanamycin. Phenotypic screening and sequence confirmation of the *nifHDK* site (using appropriate primers; **Table S3**) was performed as described above.

### Fluorescence knockdown assay

Fluorescence assays followed Banta et al.^36^ with modifications as described below. Seed cultures (nine biological replicates per strain) were inoculated from solid BN medium plates and grown in 50 mL liquid BN medium for 24 h. 1:100 dilutions of each seed culture were prepared in 5 mL BN medium followed by subsequent 1:100 dilutions in 50 mL BN medium + IPTG. The final dilutions were incubated for 10 doublings (24 h). 1.5 mL of each culture was then pelleted and resuspended in 1.5 mL 1x PBS, pH 7.4. 200 µL of each PBS suspension was aliquoted across a Nunc black-walled, flat-clear-bottom 96-well plate (Thermo Scientific Cat. No. 12-566-70), with five technical replicates each. OD600 and fluorescence (584 nm excitation, 607 nm emission) was measured by a CLARIOstar Plus Microplate Reader (BMG Labtech, Ortenberg, Germany). Fluorescence was reported as relative fluorescence units (RFU) normalized to OD600 and statistical significance was assessed by an unpaired t-test.

### Diazotrophic growth analysis

Diazotrophic growth characterization of *A. vinelandii* followed Carruthers et al^62^ as described below. Seed cultures (three biological replicates per strain) were inoculated from solid BN medium plates and grown in 50 mL liquid BN medium for 24 h. Cells were then inoculated into liquid B medium (as well as B medium + IPTG for dCas9 and sgRNA induction in experiments with strains AK053 and AK054) to an OD600 of 0.05 and aliquoted across a 96-well, flat-bottom plate (Greiner Bio-One Cat. No. 655161), with three to five technical replicates each. Each plate was sealed with a Breathe-Easy gas-permeable, adhesive membrane (Diversified Biotech Cat. No. BEM-1) and incubated in a SPECTROstar Nano Microplate Reader (BMG Labtech, Ortenberg, Germany), which maintained 30 °C internal temperature and 200 rpm double orbital agitation. OD600 measurements were obtained every 30 min for 3 days. Growth parameters were calculated using the R package Growthcurver^63^ and statistical significance was assessed either by one-way ANOVA with post-hoc Tukey HSD test or by unpaired t-test.

### Acetylene reduction assay

*A. vinelandii* seed cultures (two biological replicates for WT and three biological replicates for other strains) were grown in 50 mL BN medium for 24 h. Cells were then inoculated into 100 mL BN medium + IPTG for dCas9 and sgRNA induction and grown to mid-log phase (OD600 = 0.5-0.8). Cultures were pelleted at 4255 x g, 10 min at 4 °C and resuspended in 100 mL B medium + IPTG for nitrogenase derepression and grown for 4 h. Rubber septum caps were affixed to the mouth of each flask and 25 mL of headspace was removed and replaced with an equivalent volume of acetylene gas via syringe. Cultures were subsequently shaken at 30 °C, 300 rpm for 1 h, with headspace samples obtained every 20 min. Sample were analyzed by a Nexis GC-2030 gas chromatograph (Shimadzu, Kyoto, Japan) and ethylene was quantified by a standard curve with known ethylene concentrations.

Immediately following extraction of the last headspace sample, 48 mL of each culture was pelleted at 4,255 x g, 10 min at 4 °C and resuspended with 6 mL potassium phosphate buffer. 1 mL aliquots of each resuspension were pelleted once more at 17,000 x g, 10 min and flash frozen in liquid nitrogen. Frozen pellets were stored at −80 °C prior to total protein quantification and RNA extraction (see below). Total protein was determined using the Pierce BCA Protein Assay Kit (Thermo Scientific Cat. No. 23225) according to manufacturer instructions on a CLARIOstar Plus plate reader (BMG Labtech, Ortenberg, Germany). Protein concentrations were quantified by a standard curve of known bovine serum albumin concentrations.

Acetylene reduction rates were calculated from three time points and normalized to total protein. Statistical significance was assessed by one-way ANOVA and post-hoc Tukey HSD test.

### Nitrogenase gene expression analysis

Total RNA was extracted from frozen *A. vinelandii* cell pellets prepared following the acetylene reduction assay described above. Pellets were washed in 2 mL cold 10 mM NaCl + 4 mL cold RNAprotect Bacteria Reagent (Qiagen Cat. No. 76506) and repelletted at 4,255 x g, 10 min, 4 °C. Pellets were resuspended in 200 µL TE buffer, pH 8.0 (Quality Biological Cat. No. 10128-420) with 50 mg/mL lysozyme (Ward’s Science Cat. No. 470301-618) and agitated 5 min. RNA extraction was performed on ice with the RNeasy Mini Kit (Qiagen Cat. No. 74104) according to manufacturer’s instructions with additional on-column DNase I (Ambion Cat. No. AM2222) treatment (15 min at room temperature between column wash steps). RNA concentration and quality was assessed by a NanoDrop 2000 Spectrophotometer (Thermo Fisher Scientific, Waltham, MA, USA).

RT-qPCR was performed using the GoTaq 1-Step RT-qPCR kit (Promega Cat. No. A6020) using *nifH* and 16S rRNA (reference gene) primers reported by Poza-Carrión et al.^41^ (**Table S3**). 20 µL reaction mixes containing 100 ng template RNA were added to a 96-well PCR plate (Bio-Rad Cat. No. MLL9601) and sealed with adhesive Microseal ‘B’ PCR Plate Sealing Film (Bio-Rad Cat. No. MSB1001). Each biological replicate was analyzed in duplicate together with no reverse transcriptase and no template controls. Reactions were run on a CFX Connect Real-Time PCR Detection System (Bio-Rad Cat. No. 1855201) following manufacturer recommended thermal cycling parameters for the GoTaq 1-Step RT-qPCR kit. Gene expression analysis was performed using the ΔΔCt method^64^ relative to the 16S rRNA reference gene and WT reference strain, with relative expression in terms of fold change. Statistical significance was assessed by one-way ANOVA and post-hoc Tukey HSD test.

## AUTHOR CONTRIBUTIONS

S.J.R., A.K.G., and B.K. conceptualized the study and designed the experiments. S.J.R. and A.K.G. performed experiments and curated and analyzed data with input from B.K. S.J.R., A.K.G., and B.K. wrote and edited the manuscript. B.K. performed supervision, project administration, and funding acquisition. All authors reviewed the results and approved the final version of the manuscript.

## ACKNOWLEDGEMENTS

We thank Jean-Michel Ané, Lance Seefeldt, and members of the Kaçar laboratory for valuable discussions; Amy Banta and Jason Peters for providing Mobile-CRISPRi plasmids, *E. coli* strains, and constructive comments; and Dennis Dean and Valerie Cash for providing *A. vinelandii* strains DJ and DJ2566. This work was supported in part by the Hypothesis Fund Fellowship (B.K.) and the National Aeronautics and Space Administration (NASA) (80NSSC22K0546).

## SUPPLEMENTARY INFORMATION

**Table S1.**
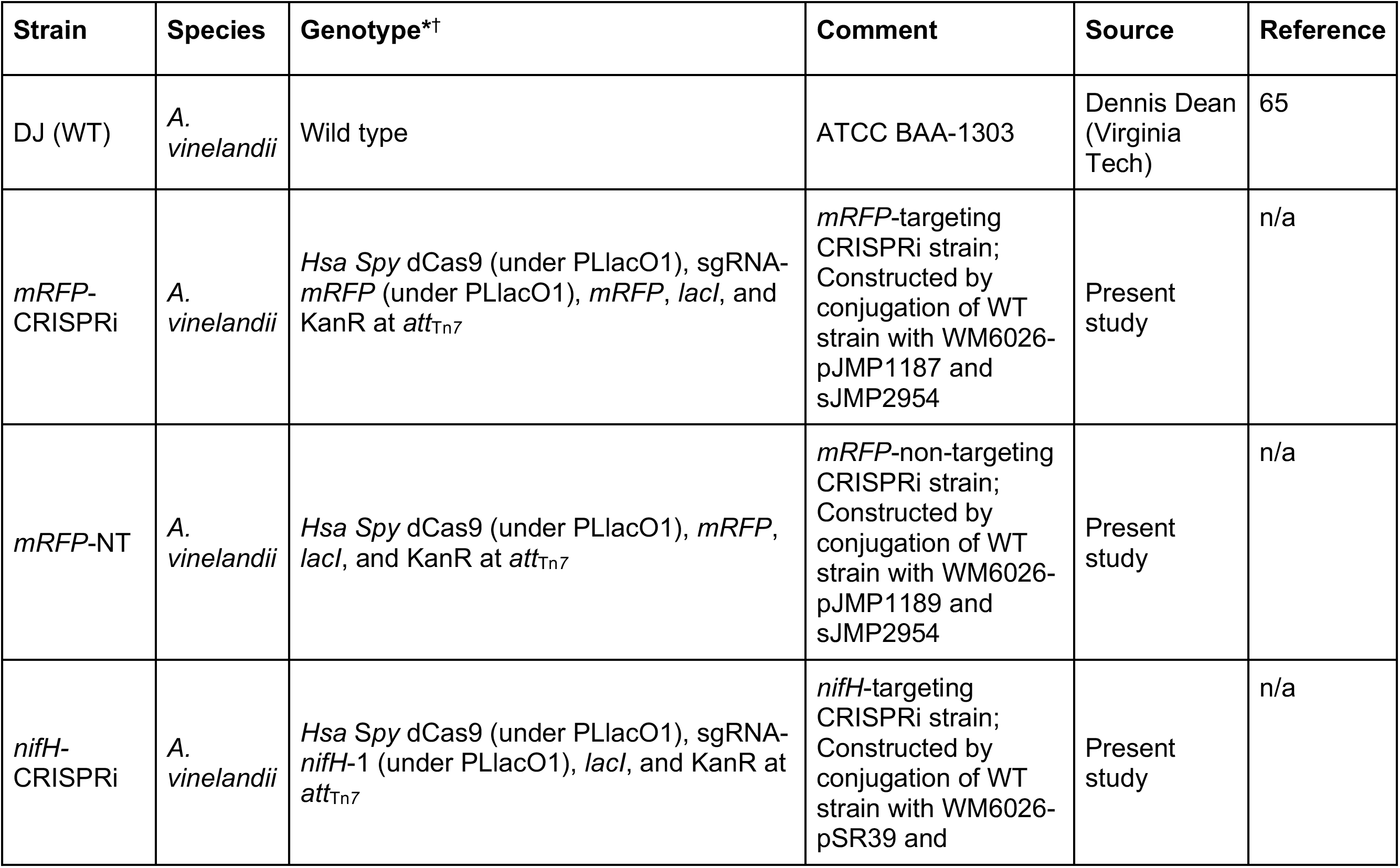
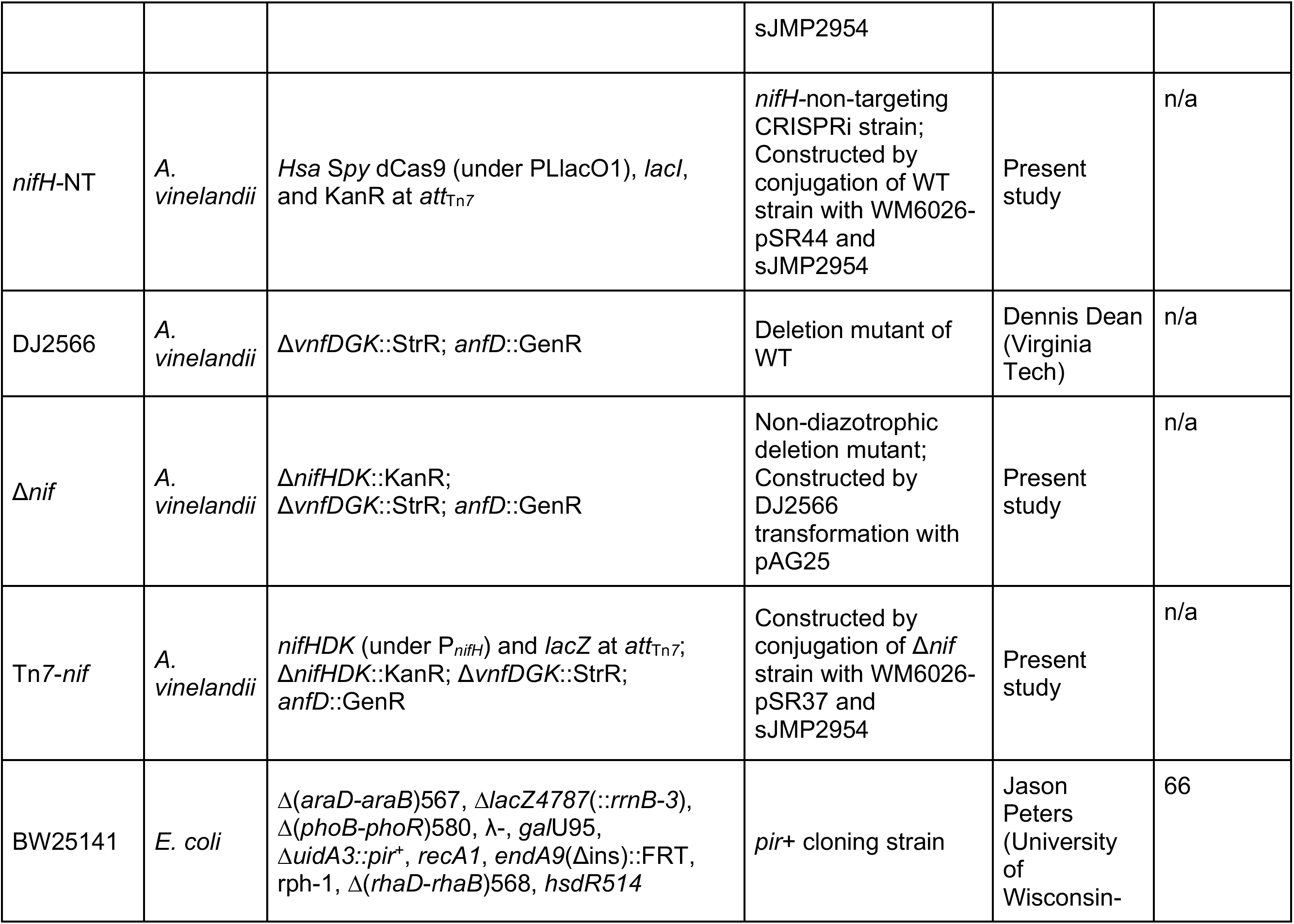
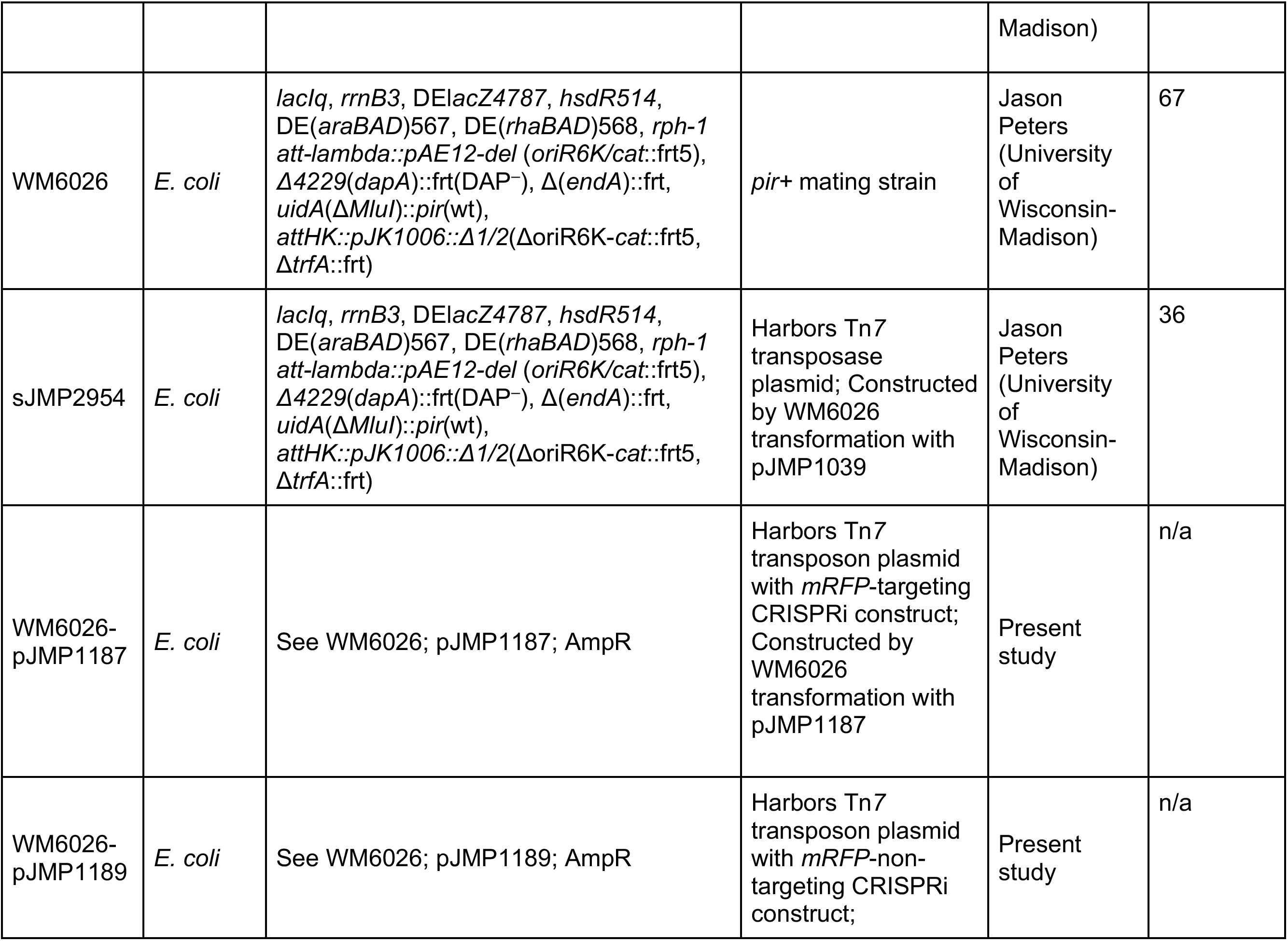
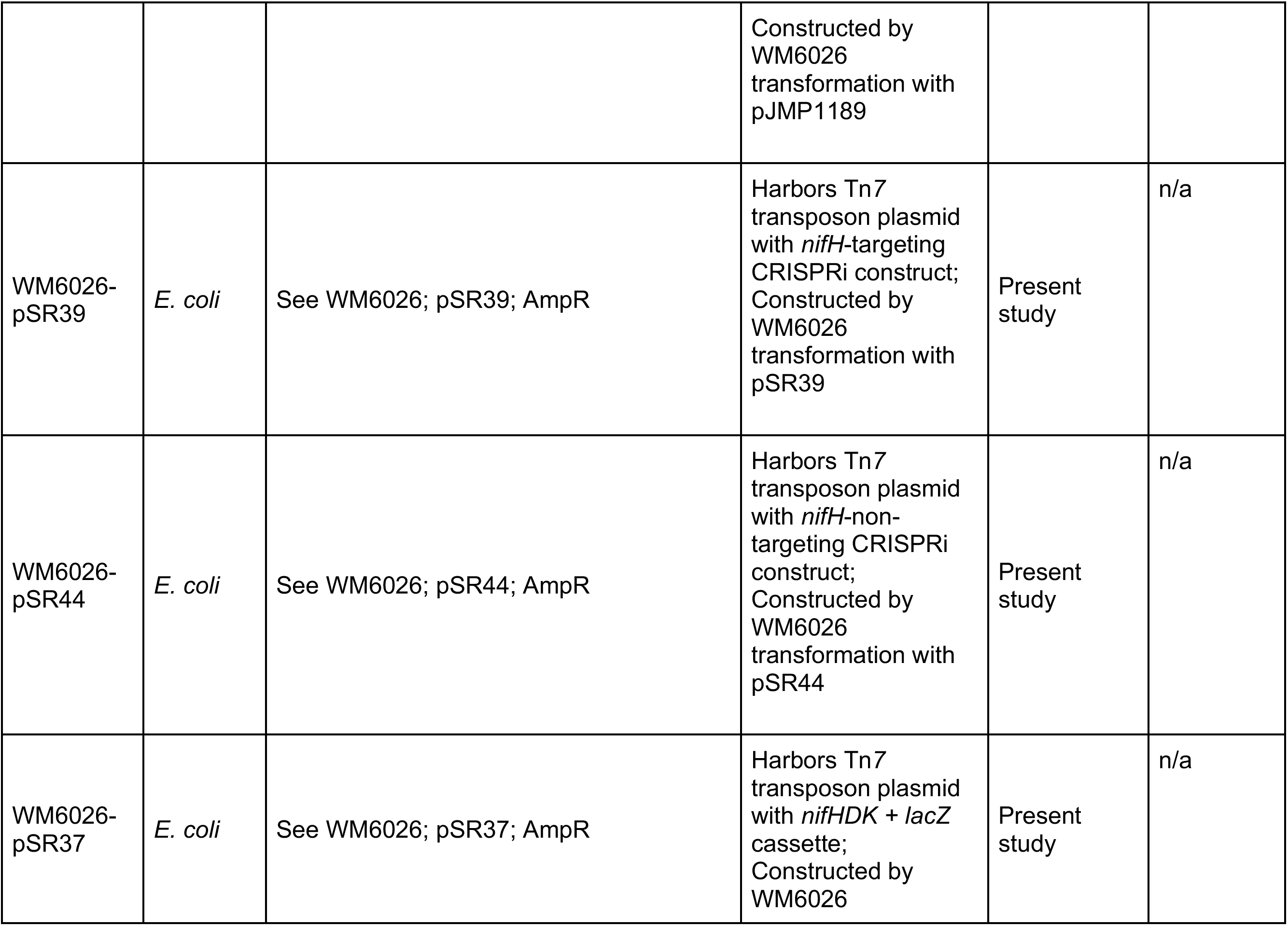
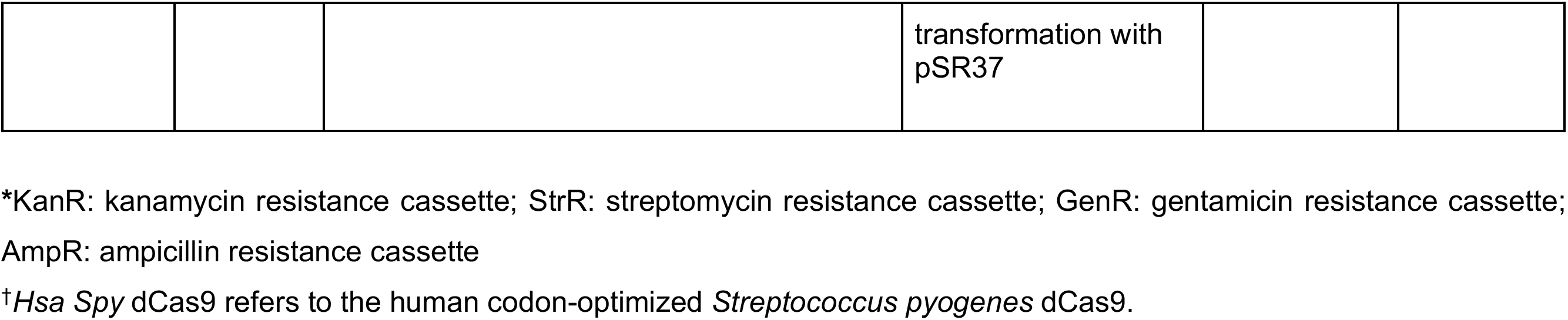
*A. vinelandii* and *E. coli* strain description and construction.

**Table S2.**
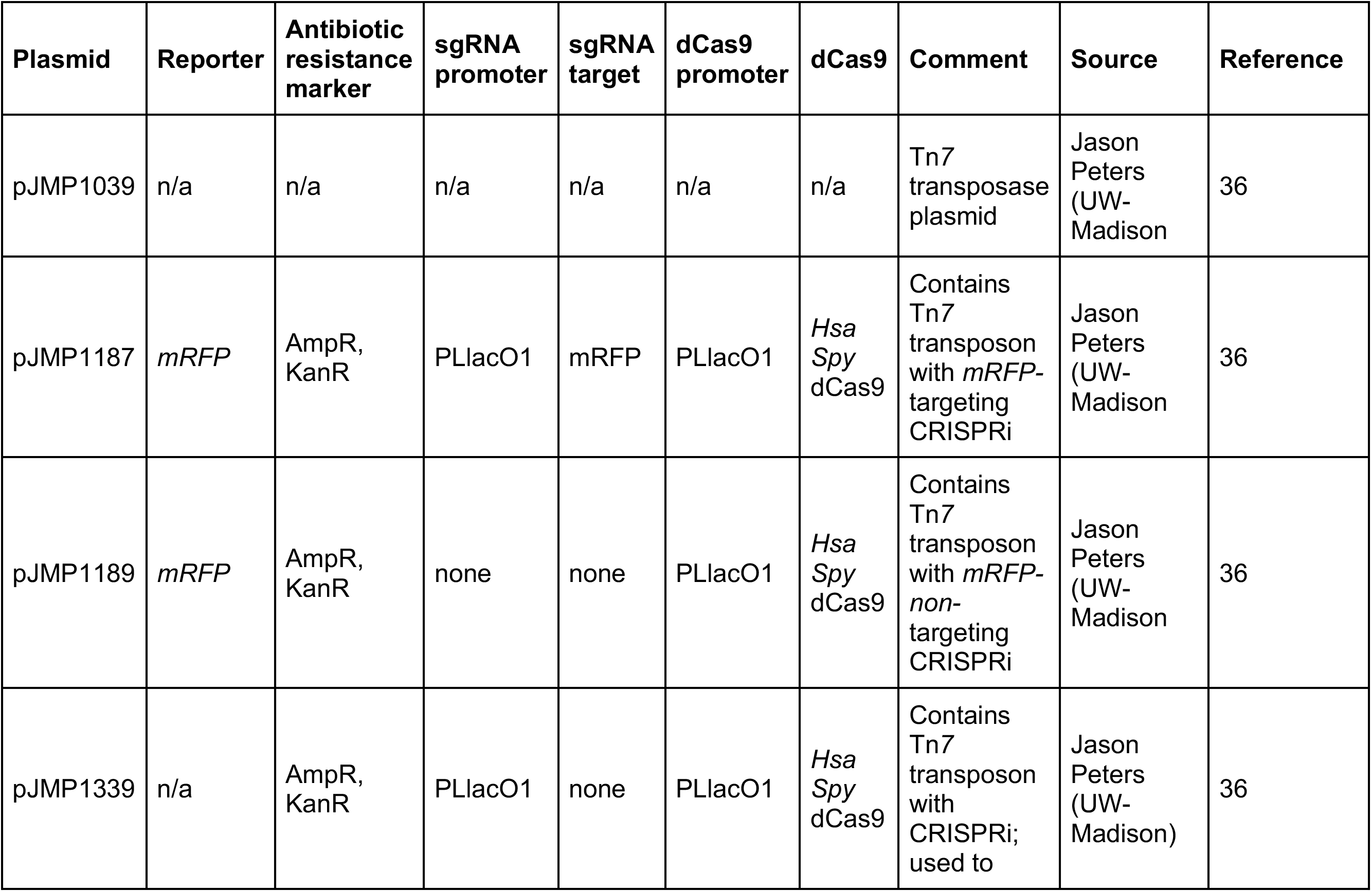
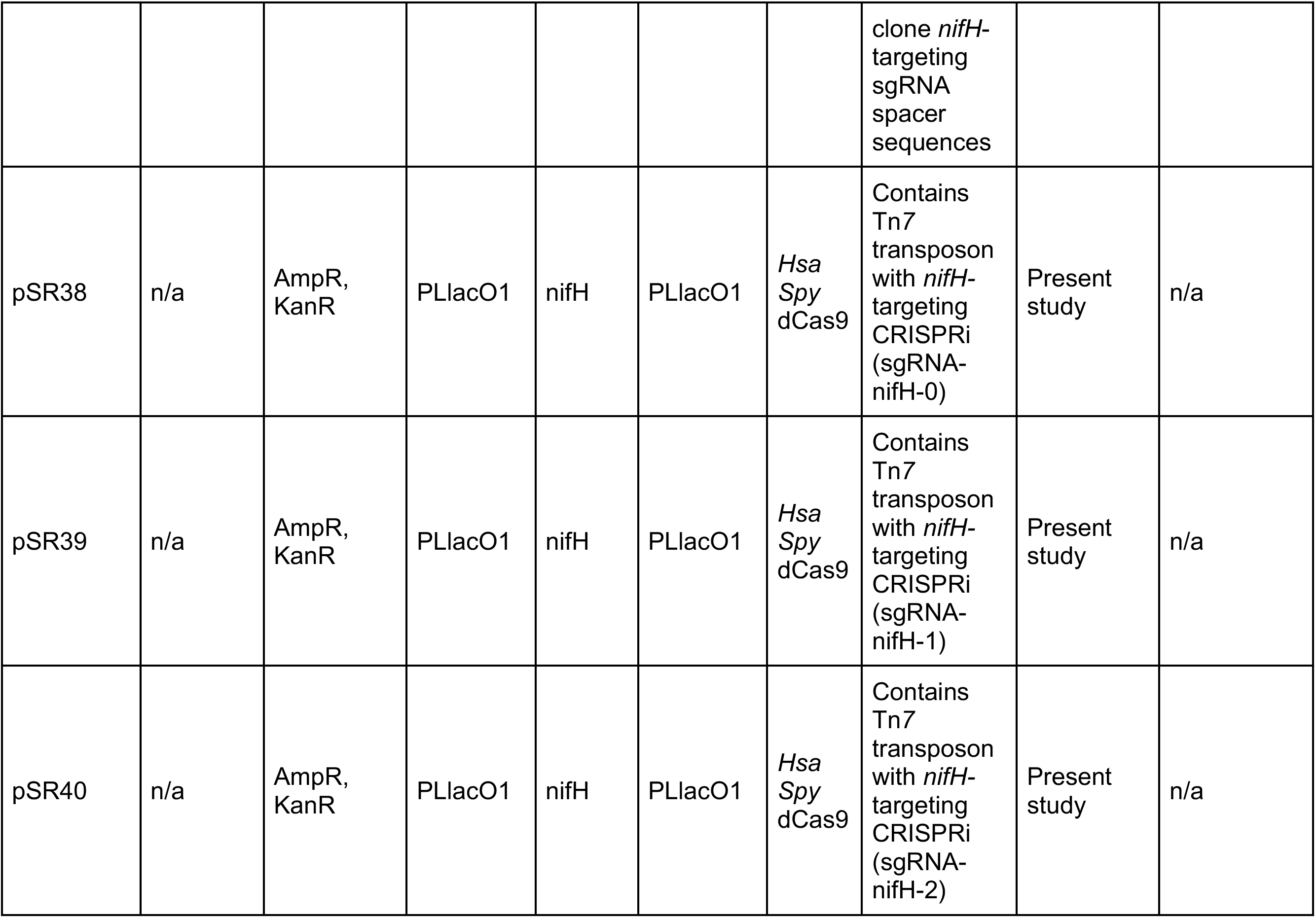
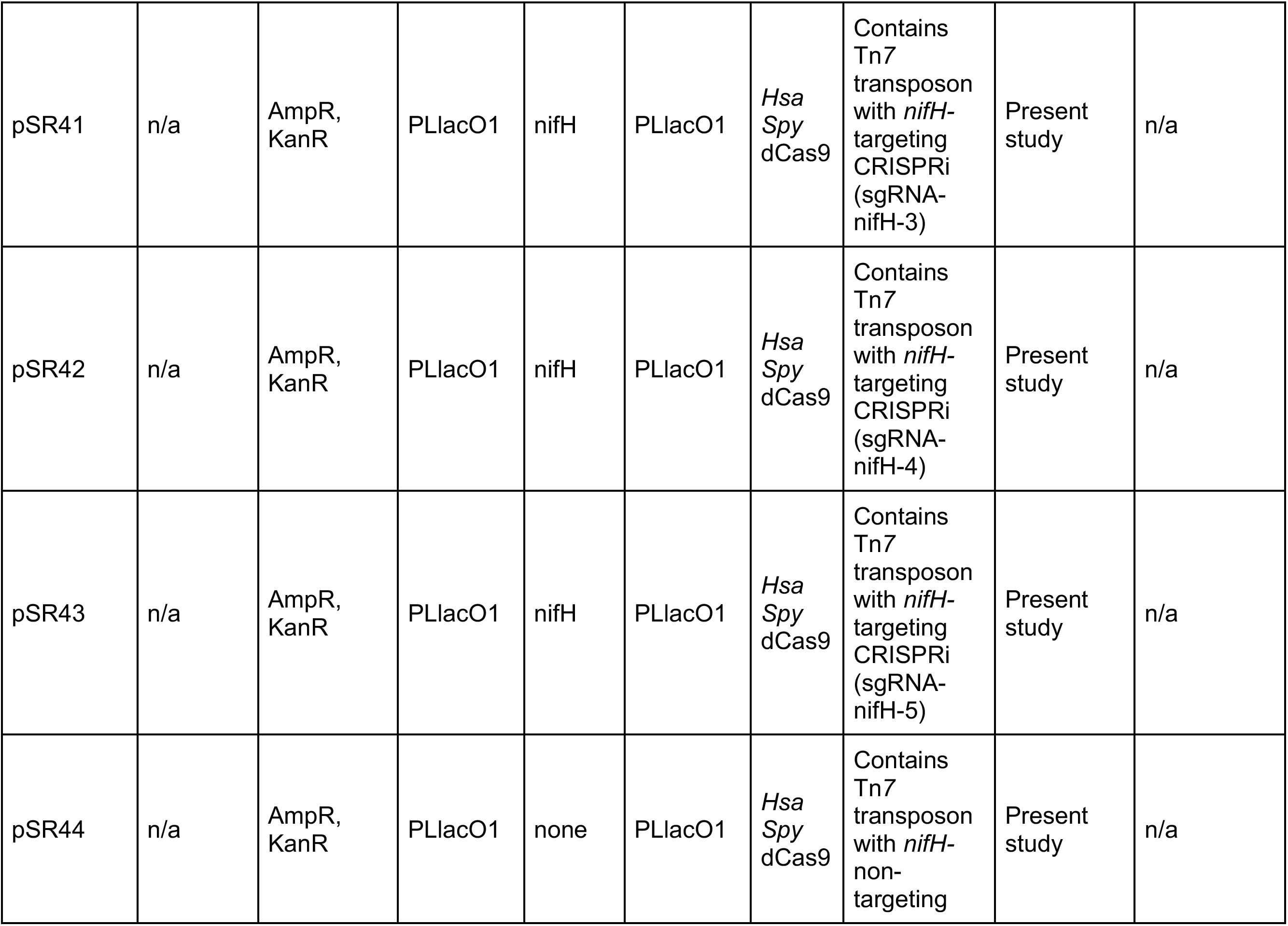
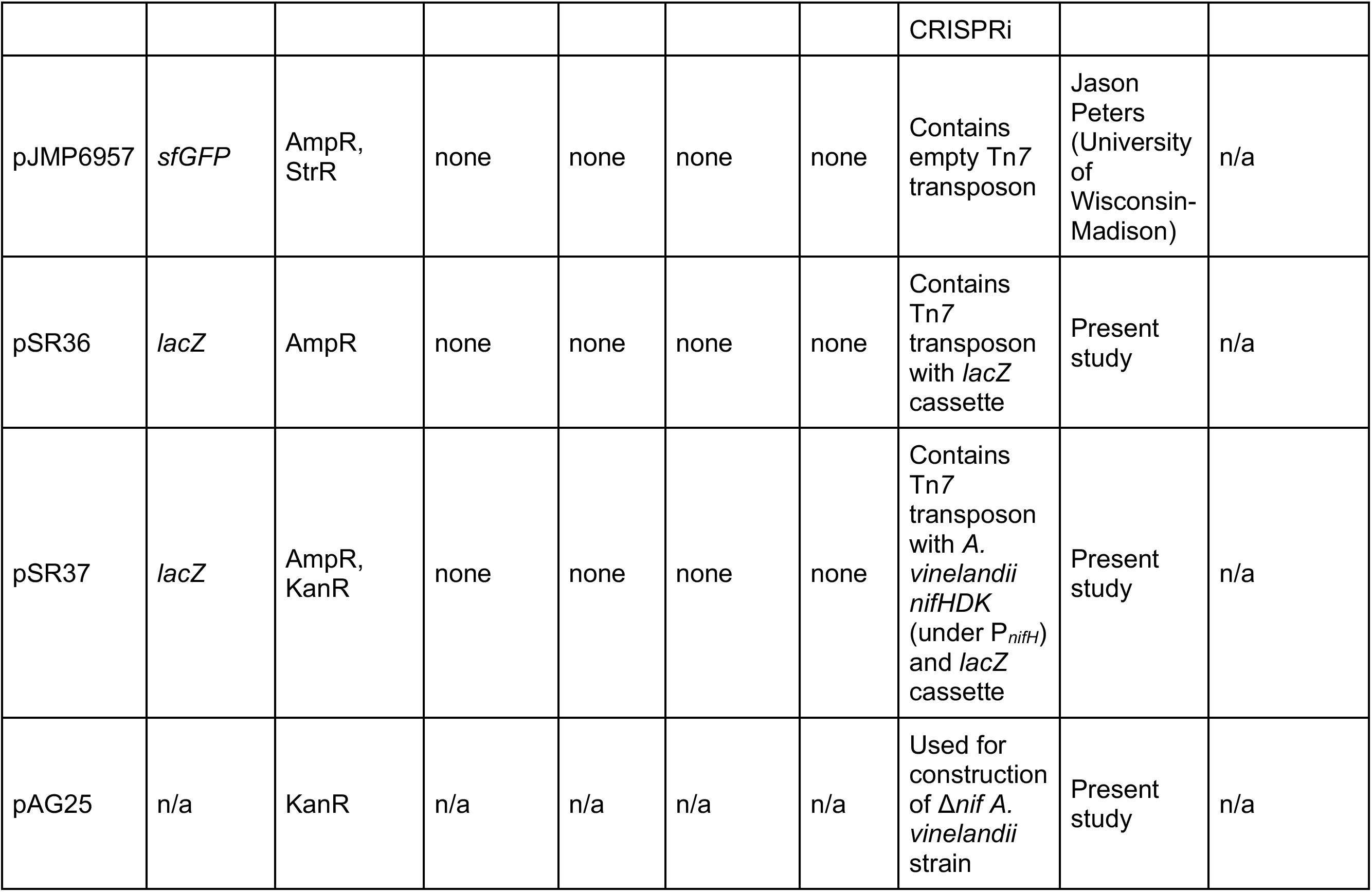
Plasmids used and constructed in the present study.

**Table S3.**
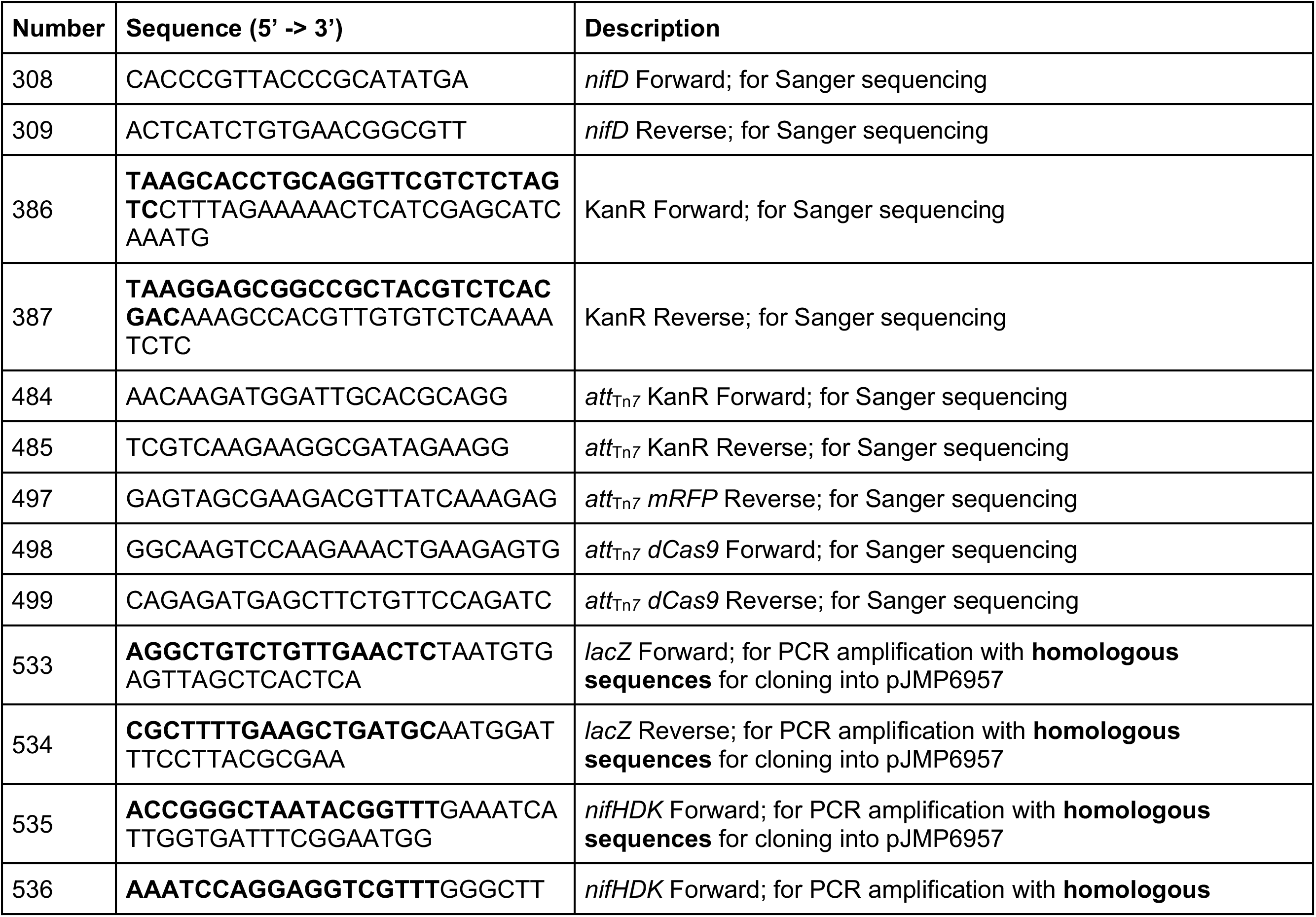
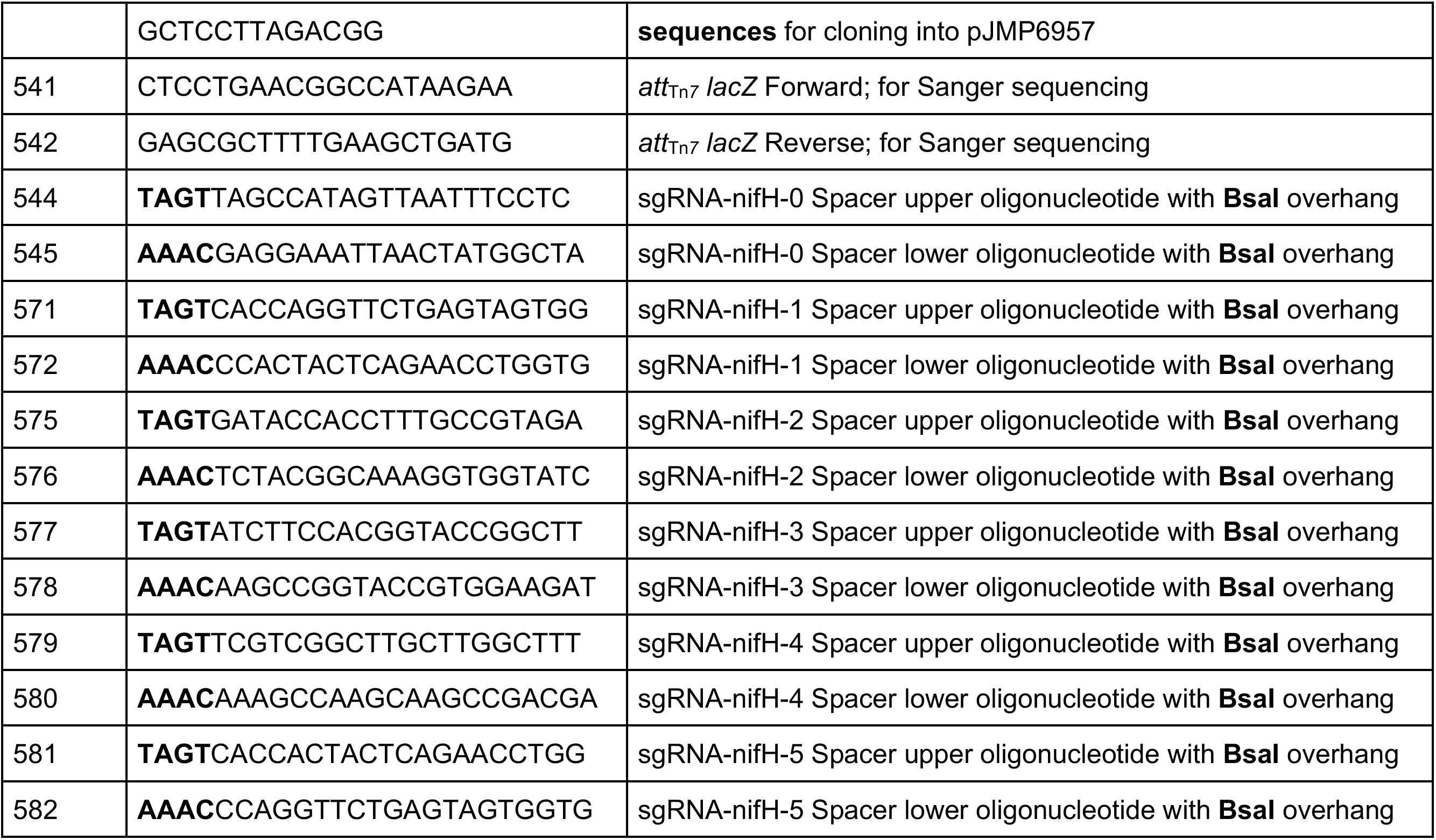
Primers and oligonucleotides generated in the present study.

**Figure S1.**
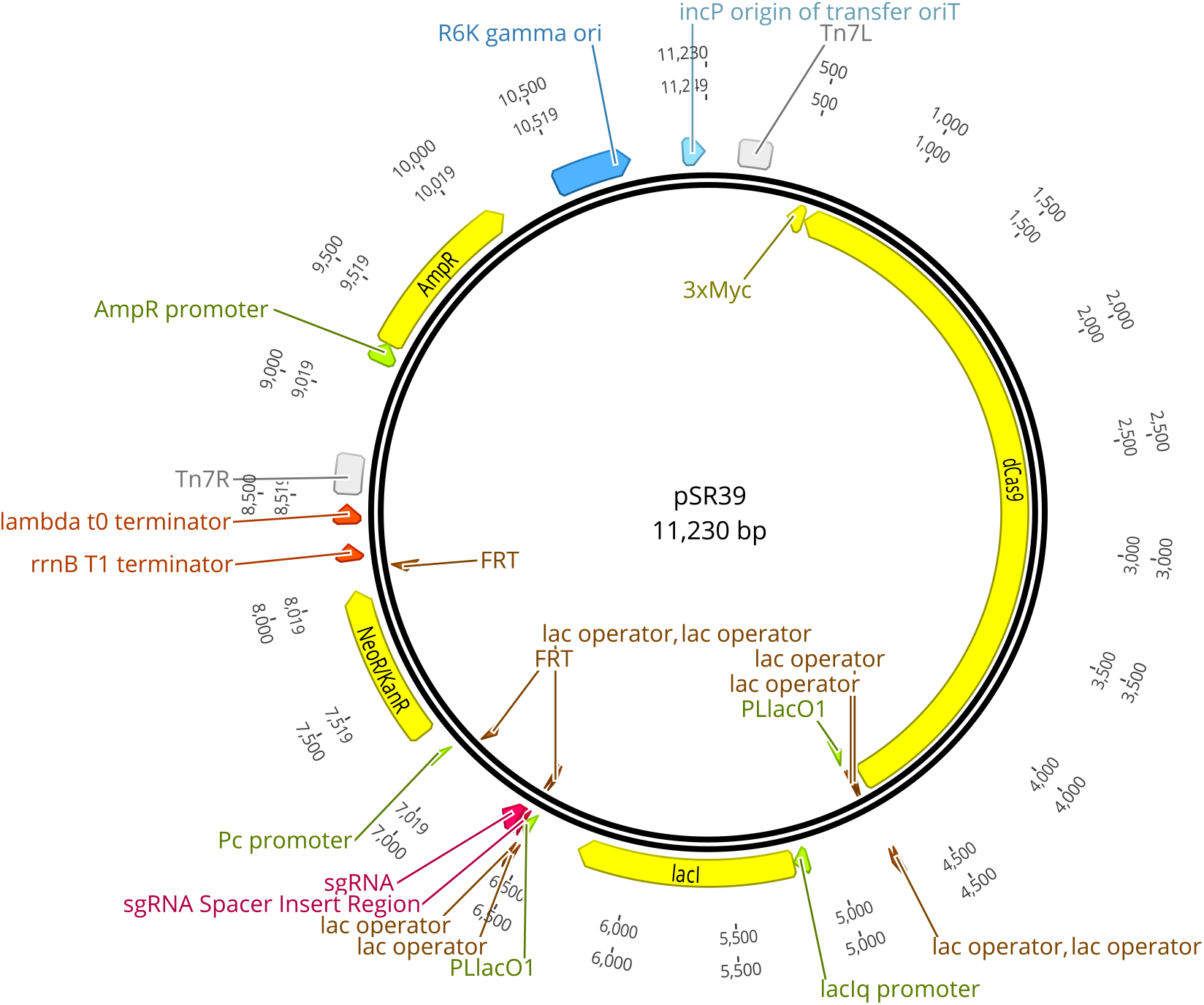
Map of plasmid pSR39, containing *nifH*-targeting CRISPRi components on a Tn*7* transposon (see **Table S2**). Map generated in Geneious Prime v2023.0.4.

**Figure S2.**
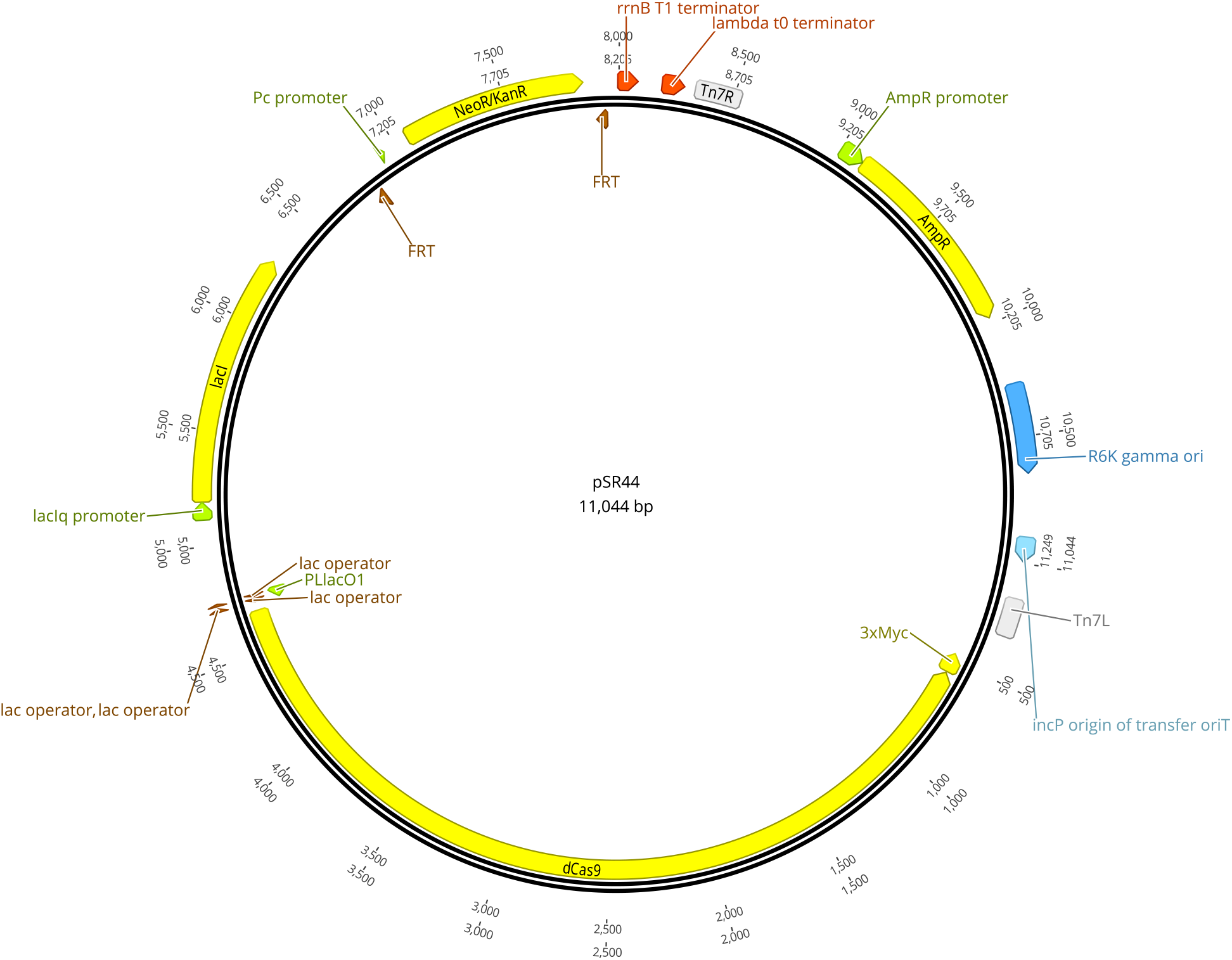
Map of plasmid pSR44, containing *nifH-non*-targeting CRISPRi components on a Tn*7* transposon (see **Table S2**). Map generated in Geneious Prime v2023.0.4.

**Figure S3.**
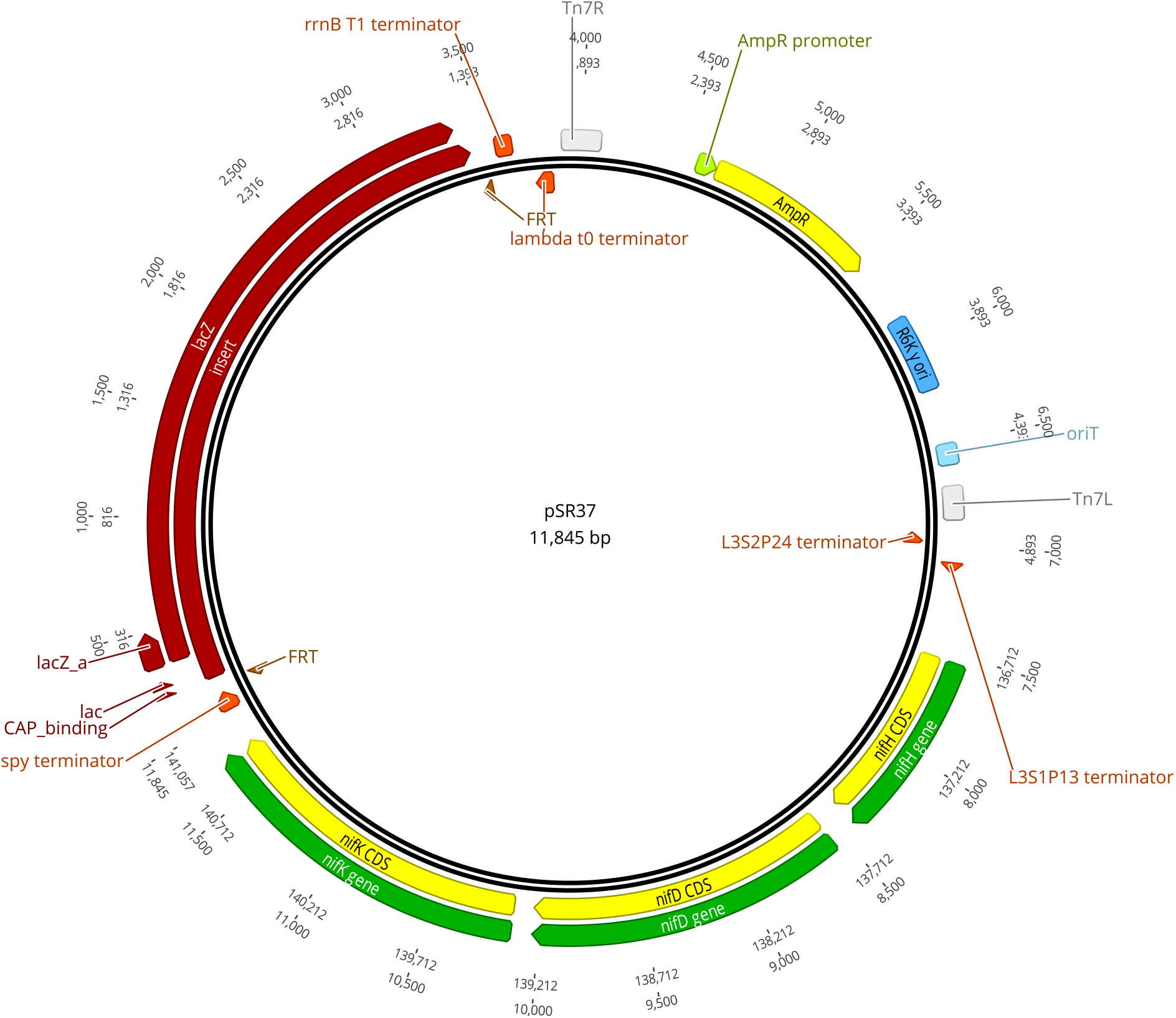
Map of plasmid pSR37, containing *nifHDK* and *lacZ* on a Tn*7* transposon (see **Table S2**). Map generated in Geneious Prime v2023.0.4.

**Figure S4.**
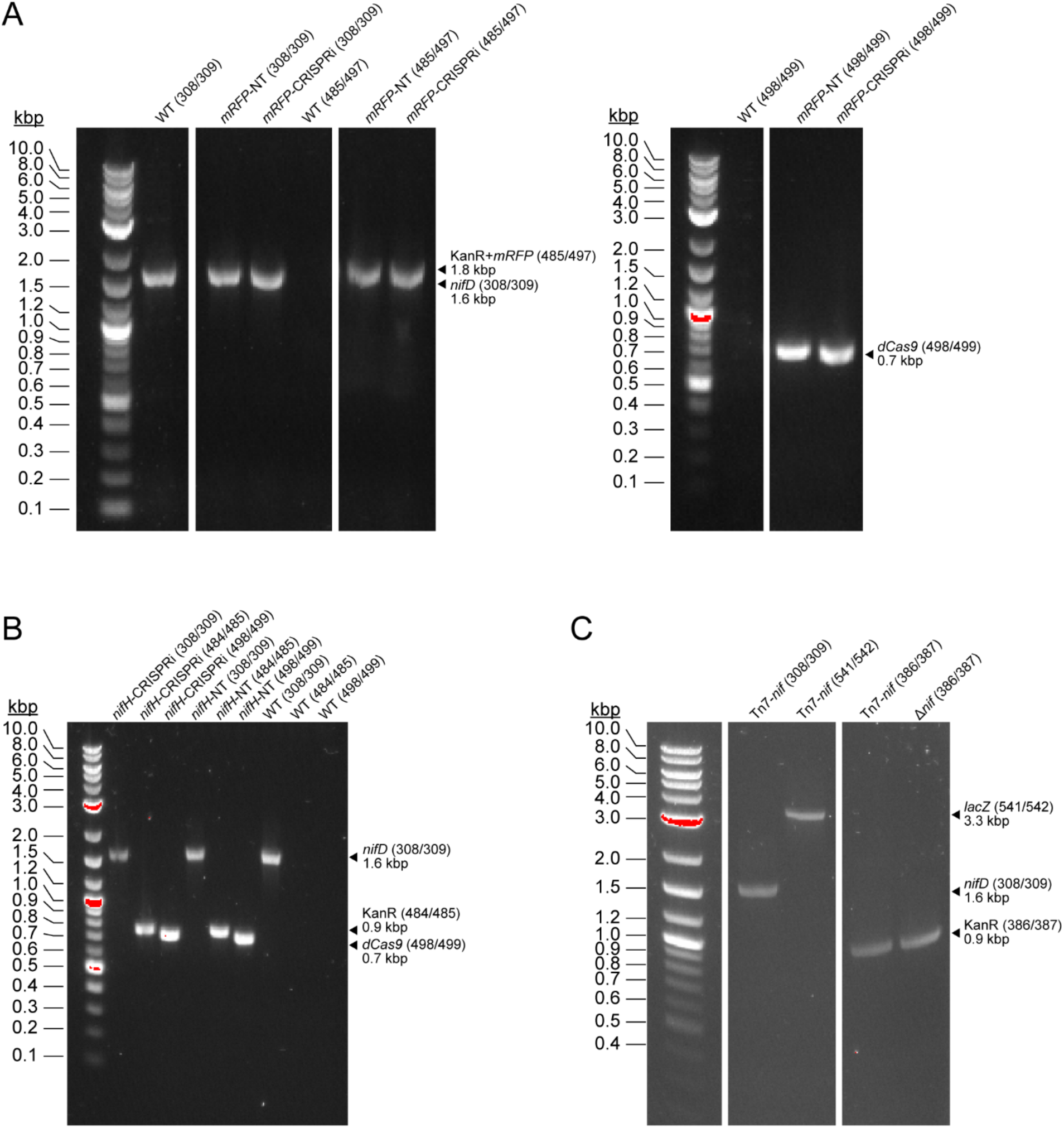
PCR-based screening of *A. vinelandii* transconjugants. PCR fragments were amplified from *A. vinelandii* gDNA (see **Methods**). (A) Screening for *mRFP*-targeting CRISPRi components inserted at *att*_Tn*7*_ (WT parent strain included as a control). Presence of native *nifD* confirms that fragments are amplified from *A. vinelandii* rather than *E. coli* donor strains. Screening is split across two gels, left and right. (B) Screening for *nifH*-targeting CRISPRi components inserted at *att*_Tn*7*_ (WT parent strain included as a control). (C) Screening for *nifHDK* and *lacZ* insertion at *att*_Tn*7*_ (Δ*nif* parent strain included as a control). (A-C) Primers used to amplify each fragment are listed in parentheses (see **Table S1** for strain descriptions and **Table S3** for primer descriptions).

## Notes

### Competing Interest Statement

The authors have declared no competing interest.

