## Supplemental Information for "A CRISPR interference system for the nitrogen-fixing bacterium *Azotobacter vinelandii*"

**SUPPLEMENTARY INFORMATION**

**Table S1.** *A. vinelandii* and *E. coli* strain description and construction.

| Strain | Species | Genotype*† | Comment | Source | Reference |
|---|---|---|---|---|---|
| DJ (WT) | *A. vinelandii* | Wild type | ATCC BAA-1303 | Dennis Dean (Virginia Tech) | 65 |
| *mRFP*-CRISPRi | *A. vinelandii* | *Hsa Spy* dCas9 (under PLlacO1), sgRNA-*mRFP* (under PLlacO1), *mRFP*, *lacI*, and KanR at $att_{Tn7}$ | *mRFP*-targeting CRISPRi strain; Constructed by conjugation of WT strain with WM6026-pJMP1187 and sJMP2954 | Present study | n/a |
| *mRFP*-NT | *A. vinelandii* | *Hsa Spy* dCas9 (under PLlacO1), *mRFP*, *lacI*, and KanR at $att_{Tn7}$ | *mRFP*-non-targeting CRISPRi strain; Constructed by conjugation of WT strain with WM6026-pJMP1189 and sJMP2954 | Present study | n/a |
| *nifH*-CRISPRi | *A. vinelandii* | *Hsa Spy* dCas9 (under PLlacO1), sgRNA-*nifH*-1 (under PLlacO1), *lacI*, and KanR at $att_{Tn7}$ | *nifH*-targeting CRISPRi strain; Constructed by conjugation of WT strain with WM6026-pSR39 and | Present study | n/a |

| | | | | | |
|---|---|---|---|---|---|
| | | | sJMP2954 | | |
| *nifH*-NT | *A. vinelandii* | *Hsa Spy* dCas9 (under PLlacO1), *lacI*, and KanR at $att_{Tn7}$ | *nifH*-non-targeting CRISPRi strain; Constructed by conjugation of WT strain with WM6026-pSR44 and sJMP2954 | Present study | n/a |
| DJ2566 | *A. vinelandii* | Δ*vnfDGK*::StrR; *anfD*::GenR | Deletion mutant of WT | Dennis Dean (Virginia Tech) | n/a |
| Δ*nif* | *A. vinelandii* | Δ*nifHDK*::KanR; Δ*vnfDGK*::StrR; *anfD*::GenR | Non-diazotrophic deletion mutant; Constructed by DJ2566 transformation with pAG25 | Present study | n/a |
| Tn*7*-*nif* | *A. vinelandii* | *nifHDK* (under $P_{nifH}$) and *lacZ* at $att_{Tn7}$; Δ*nifHDK*::KanR; Δ*vnfDGK*::StrR; *anfD*::GenR | Constructed by conjugation of Δ*nif* strain with WM6026-pSR37 and sJMP2954 | Present study | n/a |
| BW25141 | *E. coli* | Δ(*araD-araB*)567, Δ*lacZ4787*(::*rrnB-3*), Δ(*phoB-phoR*)580, λ-, *gal*U95, Δ*uidA3::pir*$^{+}$, *recA1*, *endA9*(Δins)::FRT, rph-1, Δ(*rhaD-rhaB*)568, *hsdR514* | *pir*+ cloning strain | Jason Peters (University of Wisconsin- | 66 |

| | | | | | |
|---|---|---|---|---|---|
| | | | | Madison) | |
| WM6026 | *E. coli* | *lacIq*, *rrnB3*, DElacZ4787, *hsdR514*, DE(*araBAD*)567, DE(*rhaBAD*)568, *rph-1 att-lambda::pAE12-del* (*oriR6K/cat*::frt5), *Δ4229*(*dapA*)::frt(DAP⁻), Δ(*endA*)::frt, *uidA*(Δ*MluI*)::*pir*(wt), *attHK::pJK1006::Δ1/2*(ΔoriR6K-*cat*::frt5, Δ*trfA*::frt) | *pir*+ mating strain | Jason Peters (University of Wisconsin-Madison) | 67 |
| sJMP2954 | *E. coli* | *lacIq*, *rrnB3*, DElacZ4787, *hsdR514*, DE(*araBAD*)567, DE(*rhaBAD*)568, *rph-1 att-lambda::pAE12-del* (*oriR6K/cat*::frt5), *Δ4229*(*dapA*)::frt(DAP⁻), Δ(*endA*)::frt, *uidA*(Δ*MluI*)::*pir*(wt), *attHK::pJK1006::Δ1/2*(ΔoriR6K-*cat*::frt5, Δ*trfA*::frt) | Harbors Tn*7* transposase plasmid; Constructed by WM6026 transformation with pJMP1039 | Jason Peters (University of Wisconsin-Madison) | 36 |
| WM6026-pJMP1187 | *E. coli* | See WM6026; pJMP1187; AmpR | Harbors Tn*7* transposon plasmid with *mRFP*-targeting CRISPRi construct; Constructed by WM6026 transformation with pJMP1187 | Present study | n/a |
| WM6026-pJMP1189 | *E. coli* | See WM6026; pJMP1189; AmpR | Harbors Tn*7* transposon plasmid with *mRFP*-non-targeting CRISPRi construct; | Present study | n/a |

| | | | | | |
|---|---|---|---|---|---|
| | | | Constructed by WM6026 transformation with pJMP1189 | | |
| WM6026-pSR39 | *E. coli* | See WM6026; pSR39; AmpR | Harbors Tn*7* transposon plasmid with *nifH*-targeting CRISPRi construct; Constructed by WM6026 transformation with pSR39 | Present study | n/a |
| WM6026-pSR44 | *E. coli* | See WM6026; pSR44; AmpR | Harbors Tn*7* transposon plasmid with *nifH*-non-targeting CRISPRi construct; Constructed by WM6026 transformation with pSR44 | Present study | n/a |
| WM6026-pSR37 | *E. coli* | See WM6026; pSR37; AmpR | Harbors Tn*7* transposon plasmid with *nifHDK* + *lacZ* cassette; Constructed by WM6026 | Present study | n/a |

| | | | |
|---|---|---|---|
| | | | transformation with pSR37 |

*****KanR: kanamycin resistance cassette; StrR: streptomycin resistance cassette; GenR: gentamicin resistance cassette; AmpR: ampicillin resistance cassette

†*Hsa Spy* dCas9 refers to the human codon-optimized *Streptococcus pyogenes* dCas9.

**Table S2**. Plasmids used and constructed in the present study.

| Plasmid | Reporter | Antibiotic resistance marker | sgRNA promoter | sgRNA target | dCas9 promoter | dCas9 | Comment | Source | Reference |
|---|---|---|---|---|---|---|---|---|---|
| pJMP1039 | n/a | n/a | n/a | n/a | n/a | n/a | Tn7 transposase plasmid | Jason Peters (UW-Madison | 36 |
| pJMP1187 | *mRFP* | AmpR, KanR | PLlacO1 | mRFP | PLlacO1 | *Hsa Spy* dCas9 | Contains Tn*7* transposon with *mRFP*-targeting CRISPRi | Jason Peters (UW-Madison | 36 |
| pJMP1189 | *mRFP* | AmpR, KanR | none | none | PLlacO1 | *Hsa Spy* dCas9 | Contains Tn*7* transposon with *mRFP*-*non*-targeting CRISPRi | Jason Peters (UW-Madison | 36 |
| pJMP1339 | n/a | AmpR, KanR | PLlacO1 | none | PLlacO1 | *Hsa Spy* dCas9 | Contains Tn*7* transposon with CRISPRi; used to | Jason Peters (UW-Madison) | 36 |

| | | | | | | | | | |
|---|---|---|---|---|---|---|---|---|---|
| | | | | | | | clone *nifH*-targeting sgRNA spacer sequences | | |
| pSR38 | n/a | AmpR, KanR | PLlacO1 | nifH | PLlacO1 | *Hsa Spy* dCas9 | Contains Tn*7* transposon with *nifH*-targeting CRISPRi (sgRNA-nifH-0) | Present study | n/a |
| pSR39 | n/a | AmpR, KanR | PLlacO1 | nifH | PLlacO1 | *Hsa Spy* dCas9 | Contains Tn*7* transposon with *nifH*-targeting CRISPRi (sgRNA-nifH-1) | Present study | n/a |
| pSR40 | n/a | AmpR, KanR | PLlacO1 | nifH | PLlacO1 | *Hsa Spy* dCas9 | Contains Tn*7* transposon with *nifH*-targeting CRISPRi (sgRNA-nifH-2) | Present study | n/a |

| | | | | | | | | | |
|---|---|---|---|---|---|---|---|---|---|
| pSR41 | n/a | AmpR, KanR | PLlacO1 | nifH | PLlacO1 | *Hsa Spy* dCas9 | Contains Tn*7* transposon with *nifH*-targeting CRISPRi (sgRNA-nifH-3) | Present study | n/a |
| pSR42 | n/a | AmpR, KanR | PLlacO1 | nifH | PLlacO1 | *Hsa Spy* dCas9 | Contains Tn*7* transposon with *nifH*-targeting CRISPRi (sgRNA-nifH-4) | Present study | n/a |
| pSR43 | n/a | AmpR, KanR | PLlacO1 | nifH | PLlacO1 | *Hsa Spy* dCas9 | Contains Tn*7* transposon with *nifH*-targeting CRISPRi (sgRNA-nifH-5) | Present study | n/a |
| pSR44 | n/a | AmpR, KanR | PLlacO1 | none | PLlacO1 | *Hsa Spy* dCas9 | Contains Tn*7* transposon with *nifH*-non-targeting | Present study | n/a |

| | | | | | | | | | |
|---|---|---|---|---|---|---|---|---|---|
| | | | | | | | CRISPRi | | |
| pJMP6957 | *sfGFP* | AmpR, StrR | none | none | none | none | Contains empty Tn7 transposon | Jason Peters (University of Wisconsin-Madison) | n/a |
| pSR36 | *lacZ* | AmpR | none | none | none | none | Contains Tn*7* transposon with *lacZ* cassette | Present study | n/a |
| pSR37 | *lacZ* | AmpR, KanR | none | none | none | none | Contains Tn*7* transposon with *A. vinelandii nifHDK* (under $P_{\mathit{nifH}}$) and *lacZ* cassette | Present study | n/a |
| pAG25 | n/a | KanR | n/a | n/a | n/a | n/a | Used for construction of Δ*nif A. vinelandii* strain | Present study | n/a |

**Table S3**. Primers and oligonucleotides generated in the present study.

| Number | Sequence (5’ -> 3’) | Description |
|---|---|---|
| 308 | CACCCGTTACCCGCATATGA | *nifD* Forward; for Sanger sequencing |
| 309 | ACTCATCTGTGAACGGCGTT | *nifD* Reverse; for Sanger sequencing |
| 386 | **TAAGCACCTGCAGGTTCGTCTCTAGTC**CTTTAGAAAAACTCATCGAGCATCAAATG | KanR Forward; for Sanger sequencing |
| 387 | **TAAGGAGCGGCCGCTACGTCTCACGAC**AAAGCCACGTTGTGTCTCAAAATCTC | KanR Reverse; for Sanger sequencing |
| 484 | AACAAGATGGATTGCACGCAGG | *att*$_{\mathrm{Tn7}}$ KanR Forward; for Sanger sequencing |
| 485 | TCGTCAAGAAGGCGATAGAAGG | *att*$_{\mathrm{Tn7}}$ KanR Reverse; for Sanger sequencing |
| 497 | GAGTAGCGAAGACGTTATCAAAGAG | *att*$_{\mathrm{Tn7}}$ *mRFP* Reverse; for Sanger sequencing |
| 498 | GGCAAGTCCAAGAAACTGAAGAGTG | *att*$_{\mathrm{Tn7}}$ *dCas9* Forward; for Sanger sequencing |
| 499 | CAGAGATGAGCTTCTGTTCCAGATC | *att*$_{\mathrm{Tn7}}$ *dCas9* Reverse; for Sanger sequencing |
| 533 | **AGGCTGTCTGTTGAACTC**TAATGTGAGTTAGCTCACTCA | *lacZ* Forward; for PCR amplification with **homologous sequences** for cloning into pJMP6957 |
| 534 | **CGCTTTTGAAGCTGATGC**AATGGATTTCCTTACGCGAA | *lacZ* Reverse; for PCR amplification with **homologous sequences** for cloning into pJMP6957 |
| 535 | **ACCGGGCTAATACGGTTT**GAAATCATTGGTGATTTCGGAATGG | *nifHDK* Forward; for PCR amplification with **homologous sequences** for cloning into pJMP6957 |
| 536 | **AAATCCAGGAGGTCGTTT**GGGCTT | *nifHDK* Forward; for PCR amplification with **homologous** |

| | | |
|---|---|---|
| | GCTCCTTAGACGG | **sequences** for cloning into pJMP6957 |
| 541 | CTCCTGAACGGCCATAAGAA | $att_{Tn7}$ *lacZ* Forward; for Sanger sequencing |
| 542 | GAGCGCTTTTGAAGCTGATG | $att_{Tn7}$ *lacZ* Reverse; for Sanger sequencing |
| 544 | **TAGT**TAGCCATAGTTAATTTCCTC | sgRNA-nifH-0 Spacer upper oligonucleotide with **BsaI** overhang |
| 545 | **AAAC**GAGGAAATTAACTATGGCTA | sgRNA-nifH-0 Spacer lower oligonucleotide with **BsaI** overhang |
| 571 | **TAGT**CACCAGGTTCTGAGTAGTGG | sgRNA-nifH-1 Spacer upper oligonucleotide with **BsaI** overhang |
| 572 | **AAAC**CCACTACTCAGAACCTGGTG | sgRNA-nifH-1 Spacer lower oligonucleotide with **BsaI** overhang |
| 575 | **TAGT**GATACCACCTTTGCCGTAGA | sgRNA-nifH-2 Spacer upper oligonucleotide with **BsaI** overhang |
| 576 | **AAAC**TCTACGGCAAAGGTGGTATC | sgRNA-nifH-2 Spacer lower oligonucleotide with **BsaI** overhang |
| 577 | **TAGT**ATCTTCCACGGTACCGGCTT | sgRNA-nifH-3 Spacer upper oligonucleotide with **BsaI** overhang |
| 578 | **AAAC**AAGCCGGTACCGTGGAAGAT | sgRNA-nifH-3 Spacer lower oligonucleotide with **BsaI** overhang |
| 579 | **TAGT**TCGTCGGCTTGCTTGGCTTT | sgRNA-nifH-4 Spacer upper oligonucleotide with **BsaI** overhang |
| 580 | **AAAC**AAAGCCAAGCAAGCCGACGA | sgRNA-nifH-4 Spacer lower oligonucleotide with **BsaI** overhang |
| 581 | **TAGT**CACCACTACTCAGAACCTGG | sgRNA-nifH-5 Spacer upper oligonucleotide with **BsaI** overhang |
| 582 | **AAAC**CCAGGTTCTGAGTAGTGGTG | sgRNA-nifH-5 Spacer lower oligonucleotide with **BsaI** overhang |

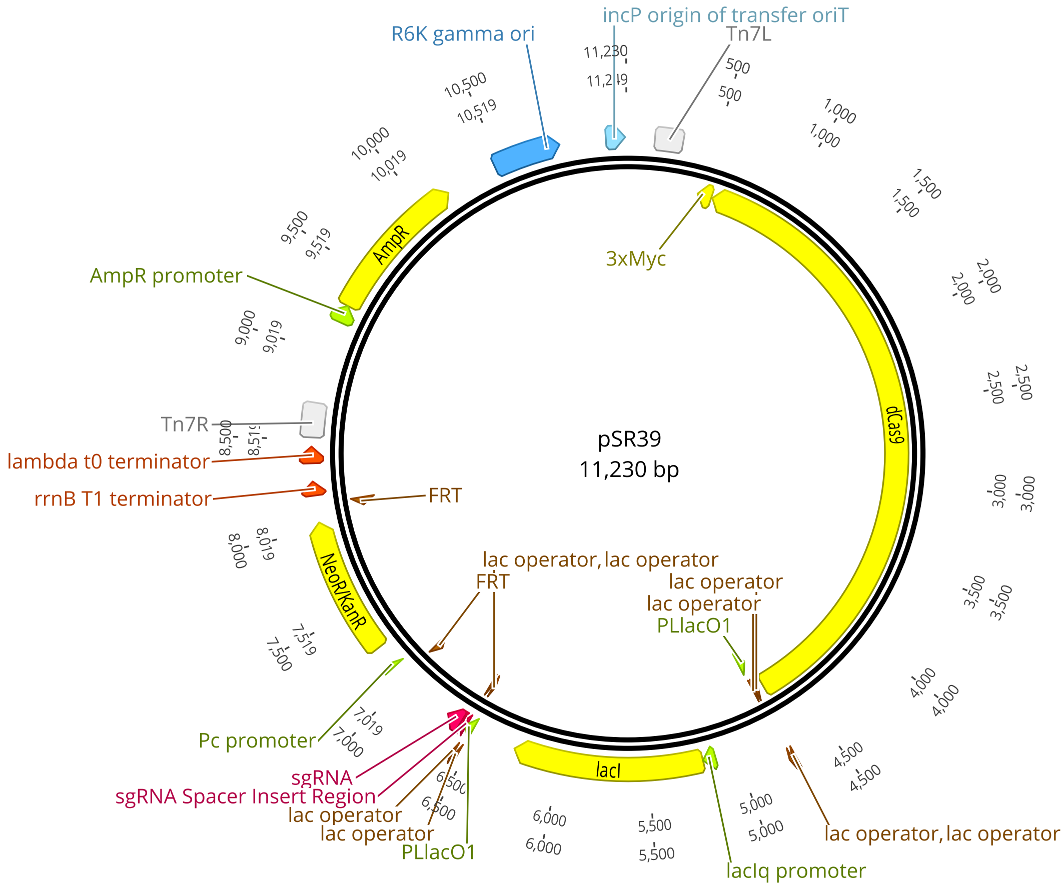

**Figure S1.** Map of plasmid pSR39, containing *nifH*-targeting CRISPRi components on a Tn*7* transposon (see **Table S2**). Map generated in Geneious Prime v2023.0.4.

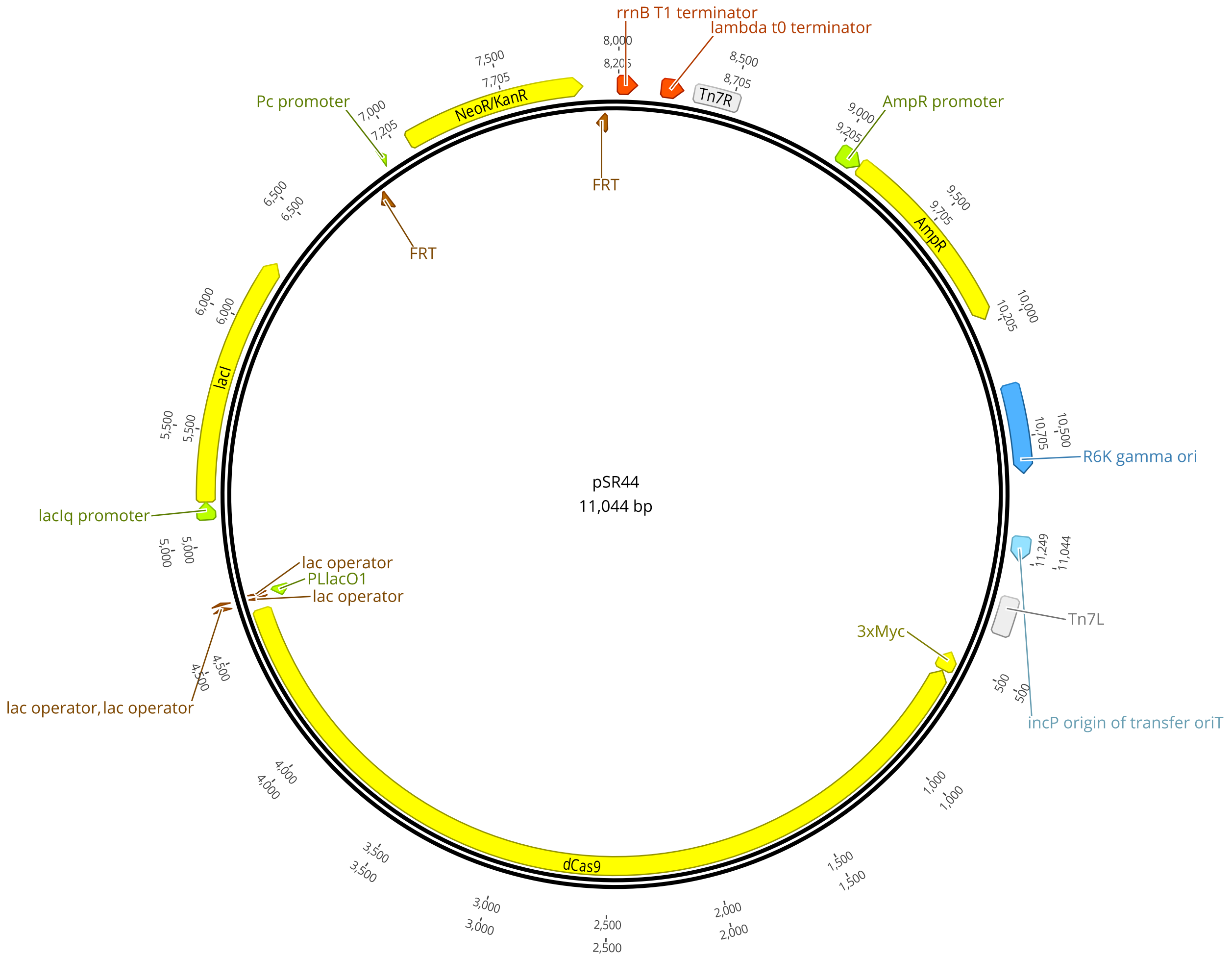

**Figure S2.** Map of plasmid pSR44, containing *nifH-non*-targeting CRISPRi components on a Tn*7* transposon (see **Table S2**). Map generated in Geneious Prime v2023.0.4.

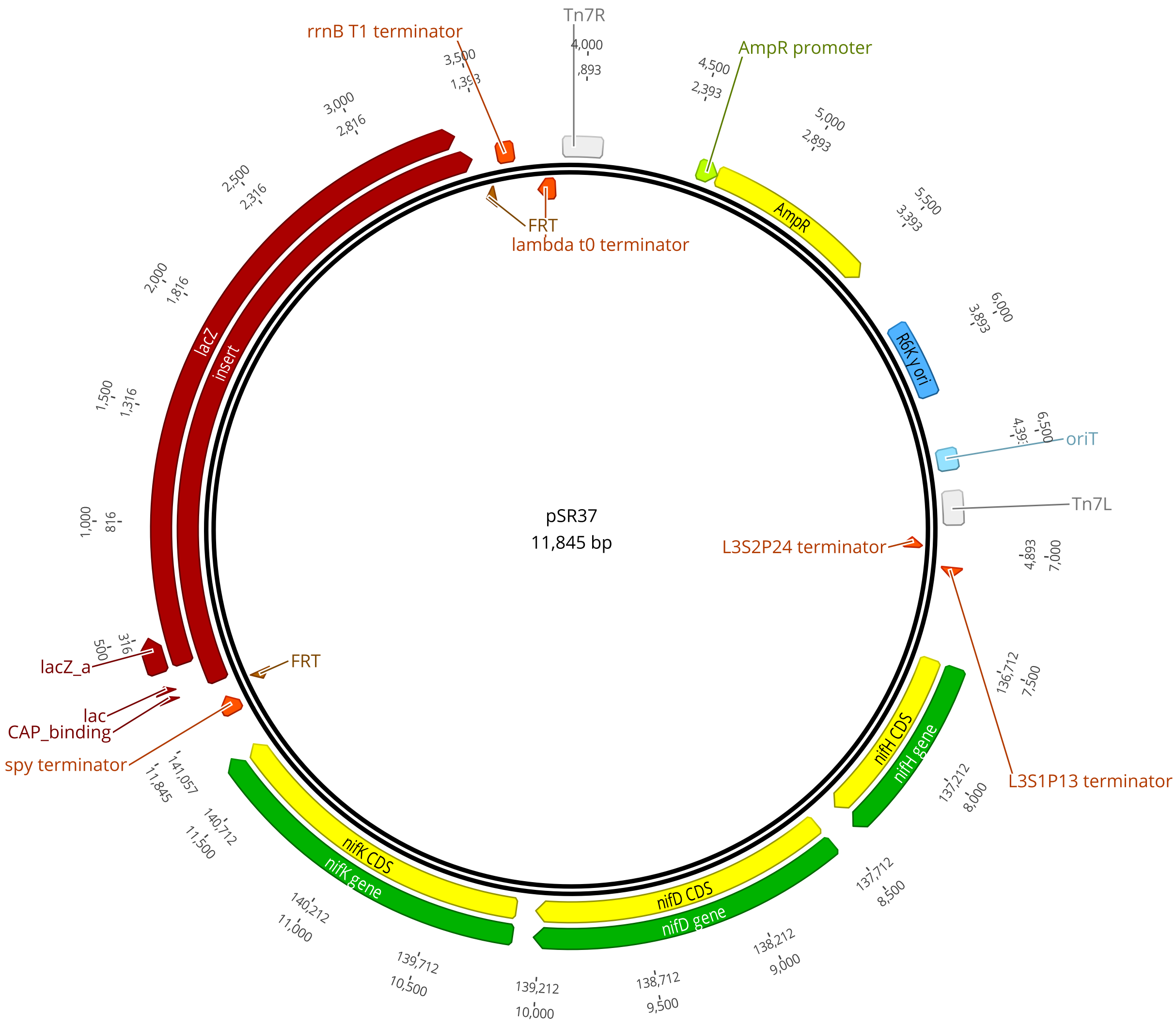

**Figure S3.** Map of plasmid pSR37, containing *nifHDK* and *lacZ* on a Tn*7* transposon (see **Table S2**). Map generated in Geneious Prime v2023.0.4.

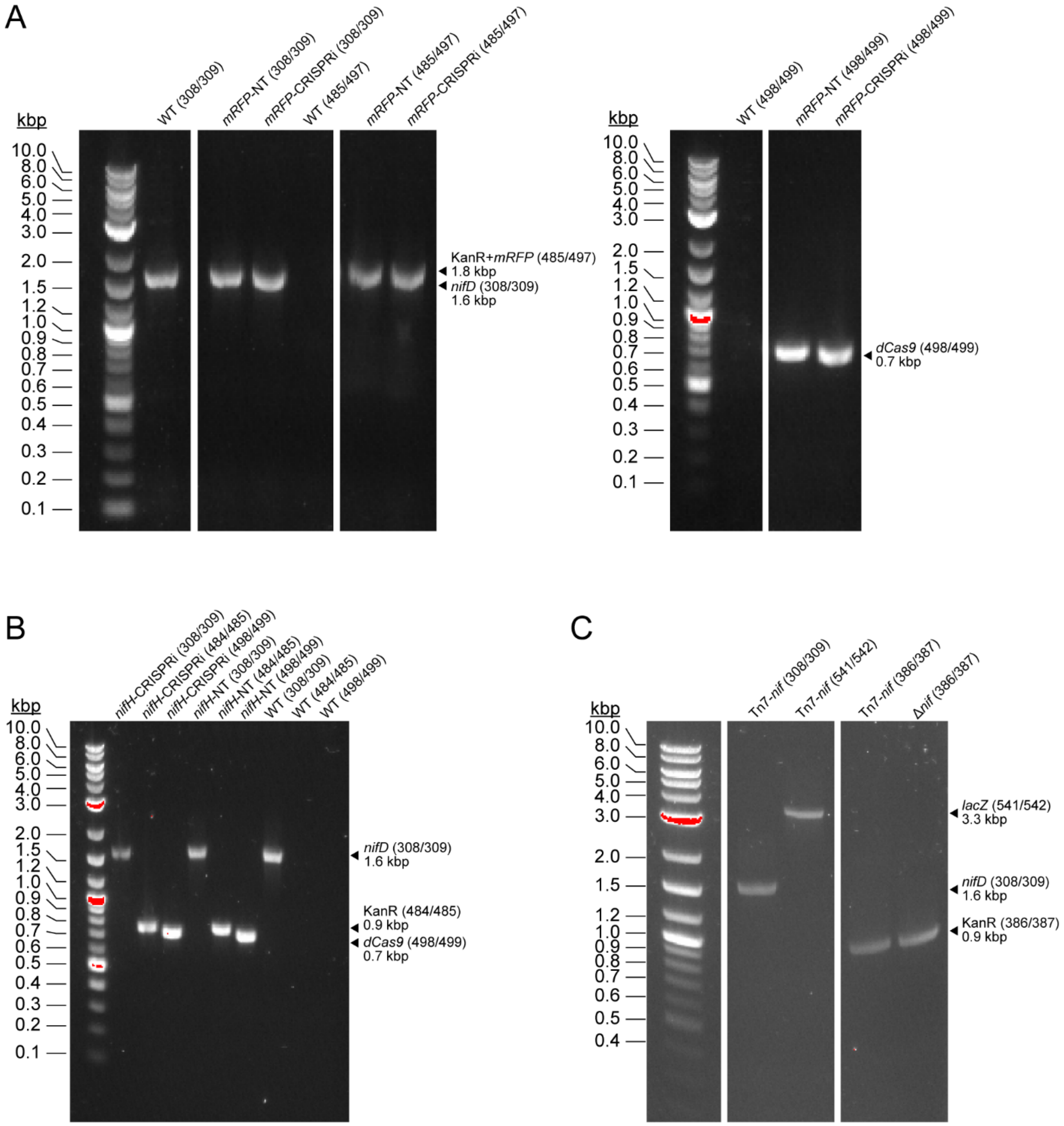

**Figure S4.** PCR-based screening of *A. vinelandii* transconjugants. PCR fragments were amplified from *A. vinelandii* gDNA (see **Methods**). (A) Screening for *mRFP*-targeting CRISPRi components inserted at *att*$_{\mathrm{Tn7}}$ (WT parent strain included as a control). Presence of native *nifD* confirms that fragments are amplified from *A. vinelandii* rather than *E. coli* donor strains. Screening is split across two gels, left and right. (B) Screening for *nifH*-

targeting CRISPRi components inserted at $att_{Tn7}$ (WT parent strain included as a control). (C) Screening for *nifHDK* and *lacZ* insertion at $att_{Tn7}$ (Δ*nif* parent strain included as a control). (A-C) Primers used to amplify each fragment are listed in parentheses (see **Table S1** for strain descriptions and **Table S3** for primer descriptions).
